# Ubiquitin recognition integrates plant cell-surface and intracellular receptor immunity

**DOI:** 10.64898/2026.02.06.703835

**Authors:** Zhishuo Wang, Tanya Mathur, Joanna Strachan, Heather Grey, Chinmayi Pednekar, Alexander von Kriegsheim, Christos Spanos, Beatriz Orosa-Puente, Steven H Spoel

## Abstract

Plant immunity is activated by cell-surface and intracellular receptors that detect pathogen-derived molecules. Mutual potentiation between these two receptor types is essential for robust disease resistance, but the mechanisms underpinning integrated dual receptor immunity remain unknown. Here, we found that activation of the intracellular receptor, ZAR1, induces its site-specific modification by non-proteolytic ubiquitin chains. ZAR1-anchored ubiquitin chains promote oligomerization of ZAR1 into a calcium-permeable resistosome pore and are recognized by RH3, a ubiquitin-binding DEAD-box RNA helicase. Remarkably, RH3 recruits both ZAR1 resistosomes and cell-surface receptor components into a dual receptor complex, thereby enhancing calcium-dependent mRNA translation of core defense proteins and establishing robust immunity. These findings identify ubiquitin recognition as the missing link in mutual potentiation of plant immunity by cell-surface and intracellular receptors.

---

Plants are continuously exposed to numerous pathogenic attackers and have evolved diverse molecular mechanisms to defend themselves. Cell surface-localized pattern recognition receptors (PRRs) detect pathogen-associated molecular patterns (PAMPs) and initiate pattern-triggered immunity (PTI). Typical PTI responses include a rapid calcium influx, reactive oxygen species (ROS) burst, activation of cytoplasmic kinase signaling, and global transcriptional and translational reprogramming to prioritize immunity over other cellular processes^1^. Adapted pathogens have evolved sophisticated strategies to subvert plant immunity by secreting effector proteins that suppress PTI. To counter this virulence strategy, plants utilize intracellular immune receptors known as nucleotide-binding domain, leucine-rich-repeat-containing receptors (NLRs) that detect pathogen effectors inside the plant cell and activate effector-triggered immunity (ETI). Upon activation, NLRs undergo conformational changes and some form oligomeric resistosome complexes that act as cation-permeable channels^2^. NLR-mediated ETI often leads to programmed cell death at the site of infection, which may restrict pathogen spread. Although PTI and ETI employ distinct signal sensors and executors, they induce highly overlapping transcriptional outputs, indicating functional synergy between these two receptor-mediated immune systems^3–5^. Indeed, physical and functional interplay between cell-surface and intracellular immune receptors has been suggested previously^6,7^. More recent studies further demonstrate mutual potentiation between cell-surface receptor-activated PTI and NLR-induced ETI^8,9^. Whereas ETI potentiates PTI-induced defense responses, cell-surface receptors are essential for optimal NLR-induced resistance outputs. This type of dual receptor signaling provides robust immunity against a wide range of adapted pathogens.

Dynamic regulation of plant immunity is critically dependent on post-translational modifications, with ubiquitination emerging as a central regulatory mechanism^10,11^. Cells modulate the stability and activity of immune proteins by covalent attachment of ubiquitin to immune receptors, immune kinases and transcription regulators^12–17^. Protein substrates can undergo monoubiquitination or polyubiquitination. In the latter, ubiquitin chains are formed by attaching the C-terminus of ubiquitin to one of eight possible residues (M1, K6, K11, K27, K29, K33, K48, K63) of the preceding ubiquitin^18^. In eukaryotes, these diverse ubiquitin chain topologies have distinct proteolytic and non-proteolytic signaling functions that are decoded by an arsenal of ubiquitin-binding proteins^19^. The majority of ubiquitin-binding proteins in plants and their functions, however, remain to be discovered. Here, we uncover numerous new ubiquitin-binding proteins and show ubiquitin recognition is the vital missing link that enables mutual potentiation of plant immunity by cell-surface and intracellular receptors.

## Ubiquitin-binding proteins are required for establishment of immunity

K48- and K63-linked ubiquitin chains represent the two most abundant topologies in eukaryotic cells, with K48 linkages often targeting substrates for proteasomal degradation, whereas K63 linkages regulate both proteolytic and non-proteolytic processes, including protein activation, complex assembly, intracellular trafficking, endocytosis, and autophagy-mediated degradation^11,20–22^. By contrast, M1-linked linear ubiquitin chains are more rare, but play a critical role in regulating immune and inflammatory signaling pathways in mammalian cells^23^. Hence, we investigated how these three ubiquitin topologies are decoded during plant immune responses. Using mass spectrometry, we profiled the global interactome of each ubiquitin topology in response to pathogen inoculation or the plant immune hormone salicylic acid (SA). Ubiquitin topology-specific pulldowns successfully captured numerous proteins that associated directly or indirectly with ubiquitin chains (Fig. 1a, Extended Data Fig. 1a, and Supplementary Data 1 and 2). While the K63 interactome was largely distinct, M1 and K48 recruited partially overlapping sets of proteins that showed Gene Ontology enrichment for immune-associated components (Fig. 1, b and c, and Extended Data Fig. 1, b and c). However, further functional annotation uncovered divergence between these two linkage types, as M1 chain-associated proteins clustered with molecular chaperones, while helicases as well as several shuttle proteins and ubiquitin receptors selectively recognized K48 chains, presumably to target unstable proteins for proteasomal degradation (Extended Data Fig. 2). By contrast, the interactome of non-proteolytic K63 chains was significantly enriched for ribosomal and other translational components, suggesting K63-binding proteins may serve as modulators of translation efficiency during immune responses (Fig. 1, b and c, Extended Data Fig. 1, b and c, and Extended Data Fig. 2, and Supplementary Data 1 and 2).

**Fig. 1.**
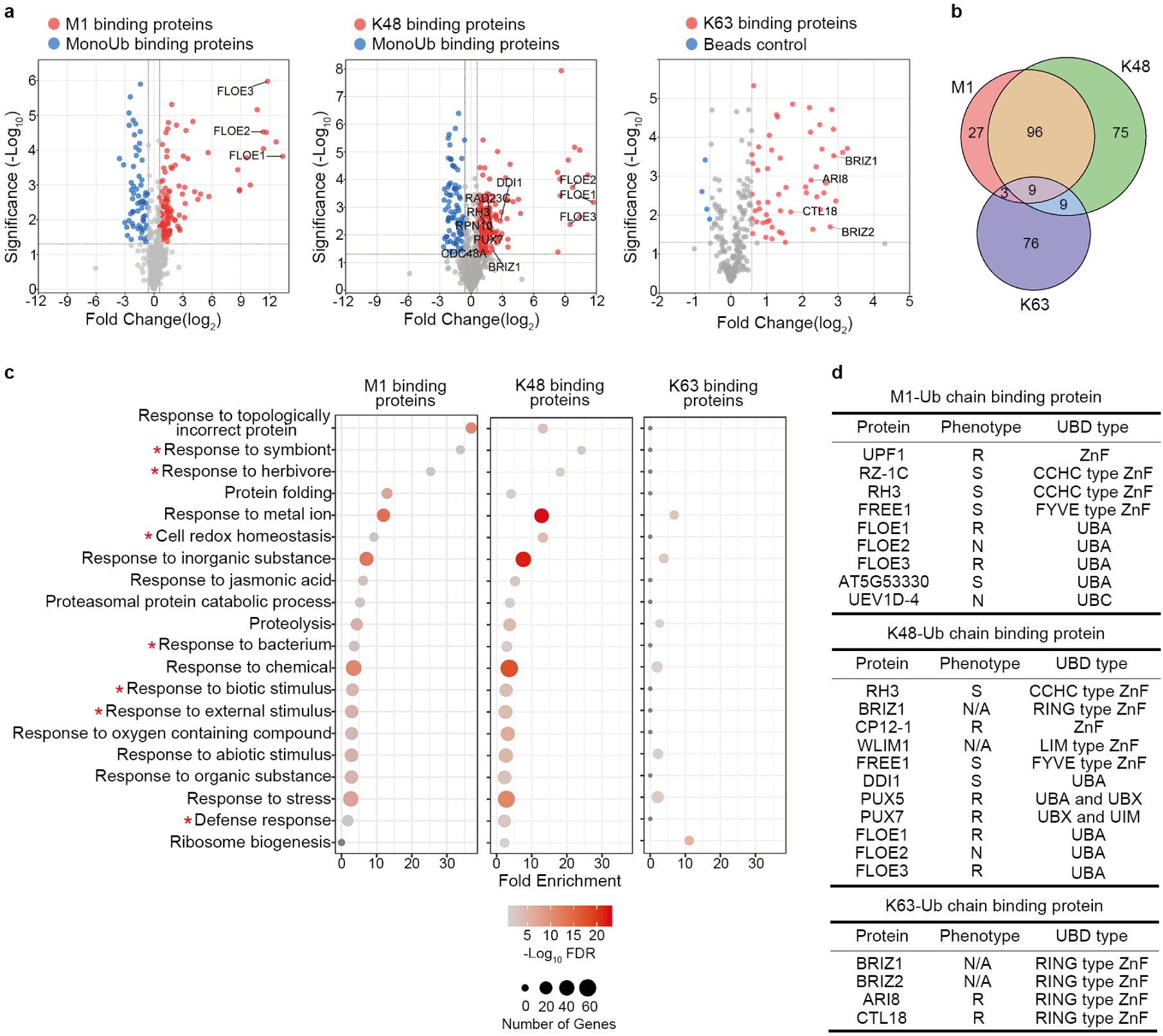
Identification of immune-responsive M1-, K48- and K63-specific ubiquitin binding proteins. **a**, Volcano plots of linkage-specific ubiquitin-binding interactomes. Adult WT plants were infected with 5 × 10^6^ CFU/mL *Psm* ES4326 for 24 h and binding proteins pulled down with indicated topologies. Monoubiquitin was used as a negative control for M1- and K48-linkage interactomes, while empty beads were used as a negative control for the K63-linkage interactome. Dashed lines indicate thresholds of 0.05 for *P*-values and 1.5 for fold changes relative to the negative control. **b**, Venn diagram showing the overlap between M1-, K48-, and K63-linked ubiquitin chain interactomes. **c**, GO term analysis of topology-specific interactomes. **d**, List of identified ubiquitin-binding domain-containing proteins and their immune response phenotypes following *Psm* ES4326 infection. Plant disease responses were classified as resistant (R), susceptible (S), or showing no significant difference (N) from the WT based on quantitative measurements of bacterial growth. N/A indicates mutants were not available for experimental analysis.

Many isolated interactors are not predicted to directly bind to polyubiquitin chains, which suggests these chains recruit and orchestrate a surprisingly wide range of cellular processes. Nonetheless, many proteins harboring well-characterized ubiquitin-binding domains were also identified, including the UBA (ubiquitin-associated domain), UBX (ubiquitin regulatory X), and UIM (ubiquitin-interacting motif) (Fig. 1d, and Supplementary Data 1 and 2). Moreover, zinc-finger (ZnF) motifs are increasingly recognized as ubiquitin-binding modules, as demonstrated by both structural and biochemical studies^19,24–26^. Our ubiquitin chain interactome analysis identified several ZnF domain-containing proteins, representing diverse ZnF protein families across all three topology-specific pulldowns (Fig. 1d)^27^. These ZnF proteins have not been previously described as ubiquitin-binding proteins *in planta*. Further *in vitro* validation confirmed that all candidate ubiquitin-binding proteins tested, including ZnF proteins, physically associated with ubiquitin chains, thus uncovering previously uncharacterized new groups of ubiquitin readers in plants (Extended Data Fig. 3a). To establish the immune functions of candidate ubiquitin-binding proteins, we analyzed disease phenotypes in corresponding mutant lines following inoculation with the virulent pathogen *Pseudomonas syringae* pv. *maculicola* (*Psm*) ES4326. Compared to the wild type (WT), the majority of mutants displayed either enhanced resistance or increased susceptibility to infection (Fig. 1d and Extended Data Fig. 3b), indicating that ubiquitin chain recognition and decoding play critical roles in orchestrating plant immune responses.

## The ubiquitin-binding protein RH3 regulates immunity induced by cell-surface receptors

Next, we explored in greater detail how ubiquitin-binding proteins may regulate plant immunity. Among the ZnF-containing ubiquitin-binding proteins, we identified DEAD-box RNA Helicase 3 (RH3), which associated with both M1- and K48-linked ubiquitin chains, but not K63-linked chains (Fig. 1d and Extended Data Fig. 3c), indicating that RH3 selectively recognizes specific ubiquitin chain topologies. RH3 and its functional domains are highly conserved across diverse plant species (Extended Data Fig. 4a). Its C-terminal CCHC-type ZnF domain is predicted to recognize diubiquitin and experimental evidence indeed verified that ZnF deletion resulted in complete loss of interaction with both M1- and K48-linked tetraubiquitin chains (Fig. 2a and Extended Data Fig. 4, b and c). RH3 was previously characterized as a chloroplast-localized helicase that functions in intron splicing^28,29^. However, in addition to its chloroplast localization, YFP-tagged RH3 also localized to the plasma membrane, occasionally forming puncta (Extended Data Fig. 4d). Membrane localization was independent of ubiquitin recognition, as mutation of the ZnF domain did not alter the subcellular distribution of RH3 (Extended Data Fig. 4d). We confirmed the subcellular localization of both endogenous RH3 and transgenic YFP-RH3 by isolating plasma membrane fractions, and found that bacterial inoculation further enhanced its abundance at the plasma membrane (Fig. 2b). Notably, an *rh3* knock-down mutant^28^ was highly susceptible to infection by *Psm* ES4326, but expression of YFP-RH3 fully complemented both the morphological and disease susceptibility phenotypes of this mutant (Fig. 2c and Extended Data Fig. 4e-g). However, expression of the ZnF-deficient YFP-RH3^ΔZnF^ mutant protein failed to restore these defects, demonstrating the indispensable role of ubiquitin recognition in RH3 immune function.

**Fig. 2.**
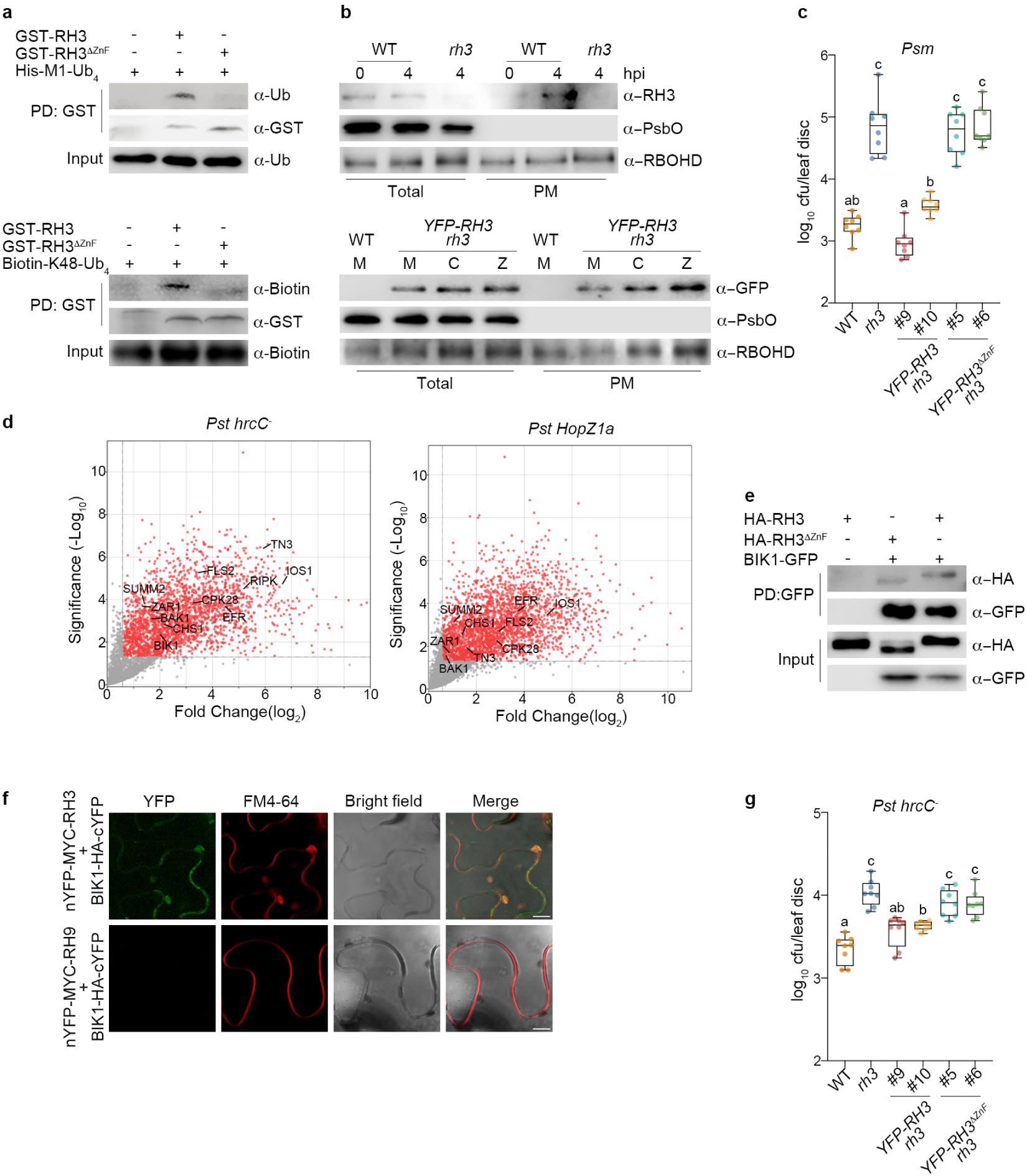
RH3 interacts with components of cell-surface receptor complexes and is required for immunity. **a**, RH3 binds both M1- and K48-linked ubiquitin chains through its ZnF domain. Recombinant GST-tagged RH3 or RH3^ΔZnF^ proteins were pulled down (PD) from *Escherichia coli* and incubated with His-tagged M1-linked tetra-ubiquitin (His-M1-Ub_4_) or biotinylated K48-linked tetra-ubiquitin (Biotin-K48-Ub_4_). His-M1-Ub_4_ and Biotin-K48-Ub_4_ were detected using anti-ubiquitin or anti-biotin, respectively. The GST fusion proteins were detected with an anti-GST antibody. **b**, Bacterial inoculation enhances RH3 accumulation at the plasma membrane. To examine the accumulation of endogenous RH3 protein, leaf samples were collected at the indicated time points after inoculation with 5 × 10^6^ CFU/mL *Pst* DC3000 (*HopZ1a*). Endogenous RH3 was detected by using an anti-RH3 antibody, PsbO and RBOHD serve as marker proteins for the chloroplast and plasma membrane, respectively. YFP-RH3 protein accumulation was assessed by immunoblot analysis using an anti-GFP antibody following inoculation with H_2_O (M), 5 × 10^6^ CFU/mL *Pst* DC3000 *hrcC⁻* (C), or 5 × 10^6^ CFU/mL *Pst* DC3000 (*HopZ1a*) (Z). **c**, The ZnF domain of RH3 is essential for establishing resistance against *Psm* ES4326. Adult plants were inoculated with 5 × 10⁵ CFU/mL bacteria and pathogen growth was measured 3 days post-inoculation (dpi). Each datapoint corresponds to a biological replicate. Box plots show median (horizontal line), 25%–75% percentiles (box) and min to max values (whiskers). Letters indicate statistically significant differences between samples (Tukey’s HSD test following one-way ANOVA; *P* < 0.05, n = 8). **d**, Volcano plots of RH3 interactors during PTI (*Pst* DC3000 *hrcC⁻*) and PTI + ETI (*Pst* DC3000 (*HopZ1a*)). Plants expressing *YFP-RH3* were inoculated with the indicated pathogen strains at a concentration of 5 × 10^6^ CFU/mL and samples were collected at 8 hpi. Protein complexes were isolated using GFP-Trap magnetic agarose. **e**, BIK1 interacts with both RH3 and RH3^ΔZnF^ *in vivo*. Proteins were transiently expressed in *Nicotiana benthamiana* and protein complexes isolated using GFP-Trap magnetic agarose, followed by immunoblot analysis with anti-GFP and anti-HA antibodies. **f**, BiFC assays revealing plasma membrane-localized interactions between BIK1 and RH3 in *N. benthamiana*. FM4-64 stains the plasma membrane. Scale bar = 10 μm. **g**, Disease susceptibility in response to *Pst* DC3000 *hrcC⁻*. Adult plants were infected with 5 × 10⁵ CFU/mL bacteria and pathogen growth was evaluated at 3 dpi as in (**c**).

To understand how RH3 regulates immunity, we employed mass spectrometry to define its interactome under different immune-activated conditions. We took advantage of the strain *P. syringae* pv. *tomato* (*Pst*) DC3000 *hrcC^−^*, which triggers only PTI because it lacks a functional type III secretion system^30,31^, and the avirulent bacterial strain *Pst* DC3000 (*HopZ1a*) that activates both PTI and HopZ-Activated Resistance 1 (ZAR1) receptor-dependent ETI^32^. The YFP-RH3 interactomes during PTI and ETI showed substantial overlap, with interactors significantly enriched in various plant defense pathways, including terms associated with PTI, ETI and systemic immunity (Fig. 2d, Extended Data Fig. 5, a and b and Supplementary Data 3). Deletion of the ZnF domain markedly diminished the RH3 interactome, highlighting the importance of ubiquitin recognition in mediating RH3–protein associations (Extended Data Fig. 5c and Supplementary Data 3). Similarly, after inoculation with *Pst* DC3000 (avrRpt2) that elicits both PTI and RPS2 receptor-dependent ETI, the YFP-RH3 interactome showed enrichment in immune-responsive pathways comparable to those observed after *Pst* DC3000 (HopZ1a) inoculation (Extended Data Fig. 5d-f and Supplementary Data 4). This suggests that RH3 may function to integrate PTI and ETI responses. Multiple core PTI components were identified as RH3 interactors (Fig. 2d, Extended Data Fig. 5d and Supplementary Data 3 and 4) and co-immunoprecipitation (co-IP) assays confirmed that *in vivo* RH3 interacts with cell-surface receptor complex components, including SOBIR1 (Suppressor of BIR1-1), and the receptor-like cytosolic kinases RIPK (RPM1-Induced Protein Kinase) and BIK1 (Botrytis-Induced Kinase 1) (Fig. 2e and Extended Data Fig. 6, a and b). We further performed a bimolecular fluorescence complementation (BiFC) assay to determine the subcellular localization of these interactions, and found that RH3 predominantly interacts with BIK1 at the plasma membrane (Fig. 2f and Extended Data Fig. 6c-e). In contrast, BIK1 did not interact with the closely related RH9 in the BiFC assay, supporting the specificity of the RH3–BIK1 interaction (Fig. 2f). However, interaction of RH3 with BIK1, SOBIR1 and RIPK was largely independent of its ZnF domain (Fig. 2e and Extended Data Fig. 6b), indicating that recruitment of RH3 to PTI components is distinct from its ZnF domain-mediated function in ubiquitin recognition. We then examined if RH3 regulates PTI. Upon inoculation with PTI-inducing *Pst* DC3000 *hrcC⁻*, the *rh3* mutant exhibited enhanced susceptibility, which was rescued by expression of *YFP-RH3* (Fig. 2g). In contrast, expression of the mutant YFP-RH3^ΔZnF^ protein failed to complement the *rh3* mutant phenotype. Thus, RH3 associates with cell surface-localized immune receptor complexes, where its function in ubiquitin recognition is essential for effective PTI responses.

## RH3 recognizes NLR-anchored ubiquitin chains to induce ETI

After inoculation with avirulent pathogen strains, the interactome of YFP-RH3 was significantly enriched for hypersensitive cell death pathways (Extended Data Fig. 5, b and f), suggesting RH3 may also regulate ETI responses. Indeed, RH3 interacted with multiple core ETI components, including NLR immune receptors and downstream signaling proteins (Fig. 2d, Extended Data Fig. 5d and Supplementary Data 3 and 4). Among these interactors, ZAR1 is a conserved NLR receptor that recognizes a broad range of pathogen effectors^33,34^. Upon activation, ZAR1 forms a calcium-permeable channel at the plasma membrane, thereby triggering robust immune responses^32,35,36^. Co-IP assays verified association between YFP-RH3 and ZAR1-HA *in vivo*, and this interaction was predominantly localized at the plasma membrane and independent of the ZAR1-guarded pseudokinase HOPZ-ETI DEFICIENT 1 (ZED1) (Fig. 3, a and b, and Extended Data Fig. 6f-i). Strikingly, deletion of RH3’s ZnF domain completely abolished interaction with ZAR1 (Fig. 3a), suggesting ubiquitin recognition is critical for association with ZAR1. Therefore, we asked if ubiquitination of ZAR1 is required for this interaction. In mapping the plant ubiquitome, we and others found that ZAR1 is ubiquitinated (Supplementary Data 5)^37^. Analyses of *P. syringae*-induced ubiquitination sites indicated that ZAR is modified at K524 (Supplementary Data 5), a highly conserved residue located near the N-terminus of its LRR domain^33^. Structural modelling revealed that K524 is positioned at an exposed surface of the oligomeric ZAR1 resistosome, suggesting it is available for ubiquitin chain conjugation (Fig. 3c). Indeed, presence of the *P. syringae* effector HopZ1a, which is directly recognized by the ZAR1 receptor complex^32^, dramatically induced total, M1- and K48-linked ubiquitination of ZAR1 *in planta* (Fig. 3d and Extended Data Fig. 7a). Interestingly, ZED1 was not required for ubiquitination of ZAR1 (Extended Data Fig. 7b), probably because detection of transiently expressed HopZ1a by NLR receptors of *Nicotiana benthamiana* is sufficient to trigger ubiquitination of Arabidopsis ZAR1. Replacement of K524 with arginine largely abolished ubiquitination (Fig. 3d and Extended Data Fig. 7a), demonstrating this particular lysine residue is the predominant site for effector-induced ubiquitin modification likely by heterotypic M1- and K48-linked chains.

**Fig. 3.**
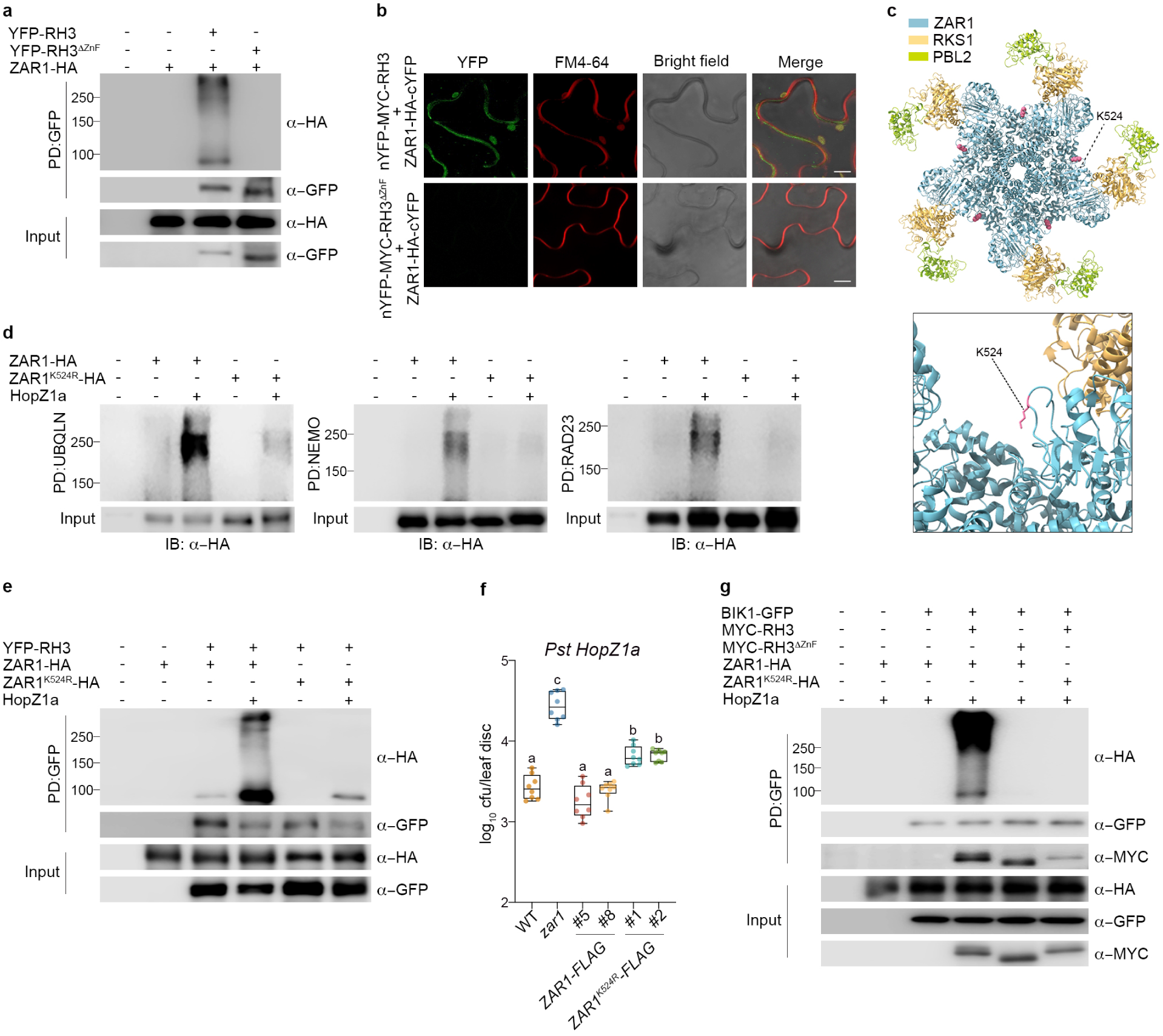
RH3 assembles ubiquitinated ZAR1 and cell-surface receptor components into a dual receptor complex. **a**, The ZnF domain of RH3 is indispensable for its interaction with ZAR1 *in vivo*. Protein complexes co-expressed in *N. benthamiana* were immunoprecipitated using GFP-Trap magnetic agarose, followed by immunoblotting with indicated antibodies. **b**, BiFC visualization of RH3-ZAR1 interaction at the plasma membrane in *N. benthamiana*. FM4-64 stains the plasma membrane. Scale bar = 10 μm. **c**, The K524 residue is located on an exposed surface of the ZAR1 resistosome. **d**, Bacterial effector HopZ1a induces ZAR1 ubiquitination with M1- and K48-linked chains at K524. Total, M1-, and K48-linked ubiquitinated substrates in *N. benthamiana* were purified using recombinant Ubiquilin, NEMO, and RAD23, respectively. ZAR1-HA ubiquitination was detected by immunoblotting with an anti-HA antibody. **e**, The K524R substitution in ZAR1 abolishes its interaction with RH3. Proteins were transiently expressed in *N. benthamiana*, immunoprecipitated using GFP-Trap and detected with indicated antibodies. **f**, The K524 residue is essential for ZAR1-mediated immune signaling. Plants were infected with 5 × 10⁵ CFU/mL *Pst* DC3000 (*HopZ1a*) and pathogen growth was evaluated at 3 dpi as in Fig 2c. **g**, Ubiquitination of ZAR1 promotes its association with BIK1-RH3 complex. Proteins were transiently expressed in *N. benthamiana*, co-immunoprecipitated using GFP-Trap and detected with indicated antibodies.

We then asked if ubiquitination of ZAR1 is required for its interaction with RH3? Co-IP assays revealed that the presence of HopZ1a enhanced interaction between RH3 and ZAR1, whereas mutation of K524 markedly reduced this interaction (Fig. 3e). Thus, RH3 recognizes ZAR1-anchored ubiquitin chains especially in presence of its effector target. Given that ZAR1 forms a pentameric resistosome upon activation^38^, we then investigated if ubiquitination at K524 affects its oligomerization. Consistent with previous findings^38^, ZAR1-HA formed oligomers when co-expressed with the kinase ZED1 and effector HopZ1a, whereas mutation of W150 impaired its oligomerization (Extended Data Fig. 7c). Similar to the ZAR1^W150A^ mutant, the K524R substitution reduced ZAR1 oligomerization (Extended Data Fig. 7c). Because ZAR1 has been reported to form oligomeric resistosomes *in vitro* in absence of ubiquitin^36^, these findings suggest that ubiquitination at K524 promotes resistosome assembly *in vivo*. Therefore, it is plausible that RH3 sequesters ubiquitinated ZAR1 monomers that subsequently form a resistosome. However, the presence of RH3 did not affect ZAR1 oligomerization and RH3 pulled down fewer oligomerization-deficient ZAR1^W150A^ than wild-type ZAR1 protein (Extended Data Fig. 7, d and e). Thus, ubiquitin likely acts as a molecular chaperone in ZAR1 resistosome assembly, but this function is independent of ubiquitin recognition by RH3.

To investigate if ubiquitination of ZAR1 is a necessary feature of plant immunity, we generated lines expressing ZAR1 variants driven by the ZAR1 promoter (*pZAR1:ZAR1-FLAG* and *pZAR1:ZAR1^K524R^-FLAG*) in the *zar1* mutant background. Expression of *ZAR1-FLAG* largely complemented enhanced susceptibility of the *zar1* mutant to avirulent *Pst* DC3000 (*HopZ1a*) (Fig. 3f). By contrast, expression of the ubiquitination-deficient *ZAR1^K524R^-FLAG* variant failed to rescue the immune-deficient phenotype of *zar1* (Fig. 3f). Together, these data demonstrate that RH3 recognizes site-specific ubiquitination of K524, which is essential for ZAR1-induced ETI responses.

## RH3 assembles a dual receptor complex to potentiate immunity through a translational boost

A recent study demonstrated that multiple intracellular NLRs serve as direct substrates of BIK1, with BIK1-mediated phosphorylation regulating NLR oligomerization and activation^39^. Because RH3 interacts with both cell-surface and intracellular immune receptors, we investigated if RH3 assembles a dual receptor complex. As shown earlier (Fig. 2e and Extended Data Fig. 6a), RH3 interacts with BIK1, which associates with cell-surface receptor complexes to activate PTI ^40–42^. Remarkably, we discovered that BIK1 also interacts with ZAR1 in an RH3-dependent manner, as their association was diminished in the absence of RH3 (Fig. 3g). Moreover, interaction was also abolished when utilizing the ZAR1^K524R^ mutant or the RH3^ΔZnF^ mutant (Fig. 3g), indicating that recognition of ZAR1-anchored ubiquitin chains by RH3’s ZnF domain is required for assembly of a dual receptor complex.

Why does RH3 recruit cell-surface and intracellular immune receptors into one and the same complex? We considered that RH3 may be the missing link in mutual potentiation of PTI and ETI immune responses induced by these two types of immune receptors^8,9^. ETI potentiates PTI defense responses via enhancing transcriptional reprogramming, leading to robust dual receptor immunity^8,9^. Indeed, we found that *Pst* DC3000 *hrcC⁻*-induced PTI initiated global transcriptional changes, which were substantially amplified during *Pst* DC3000 (*HopZ1a*)-triggered co-activation of PTI and ETI (Extended Data Fig. 8a-c and Supplementary Data 6). Although ETI-induced transcriptional reprogramming was partially attenuated in the *rh3* mutant, activation of ETI retained the capacity to potentiate PTI-responsive genes (Extended Data Fig. 8a-c), indicating that RH3 has a limited effect on immune-related transcriptional amplification during co-activation of PTI and ETI.

RNA helicases have been implicated in translational reprogramming during PTI^43^. RH3 harbors a DEAD-box RNA helicase domain that functions in intron splicing^28,29^, which is a critical step for proper mRNA maturation and subsequent protein translation^44,45^. The DEAD-box helicase domain contains a set of highly conserved motifs, including the functionally essential QxxR motif, which regulates RNA unwinding^46^. Expression of variants carrying alanine substitutions at residues Gln386 or Arg389 within the QxxR motif failed to restore the morphological and disease susceptibility phenotypes of the *rh3* mutant (Extended Data Fig. 4a and 8d-h), indicating that helicase activity of RH3 is required for immune activation. Because translational reprogramming is a hallmark of both PTI and ETI^47,48^, we hypothesized that RH3 may control plant immunity through translational regulation. To test this, we performed a surface sensing of translation (SUnSET) assay, which monitors protein synthesis rates by incorporating puromycin into nascent peptides^49^. Consistent with previous findings^47^, activation of PTI by inoculation with *Pst* DC3000 *hrcC⁻* induced significant global translational upregulation (Fig. 4, a and b). More importantly, co-activation of PTI and ETI by inoculation with *Pst* DC3000 (*HopZ1a*) strongly potentiated this translational boost. By contrast, however, translational activation and potentiation were not observed in the *rh3* mutant, indicating RH3 plays a key role in boosting protein translation during dual receptor immunity. Mutual potentiation of PTI and ETI establishes robust immunity via upregulation of multiple critical PTI components^9^. Thus, we investigated if RH3 affects protein accumulation of PTI components. In WT plants, infiltration with *Pst* DC3000 *hrcC⁻* induced protein accumulation of several PTI components, which were further boosted upon co-activation of both PTI and ETI by *Pst* DC3000 (*HopZ1a*) inoculation (Fig. 4c). In contrast, pathogen-induced protein accumulation of these PTI components was compromised in the *rh3* mutant. Hence, RH3 boosts the translation of critical defense proteins during activation of dual receptor immunity.

**Fig. 4.**
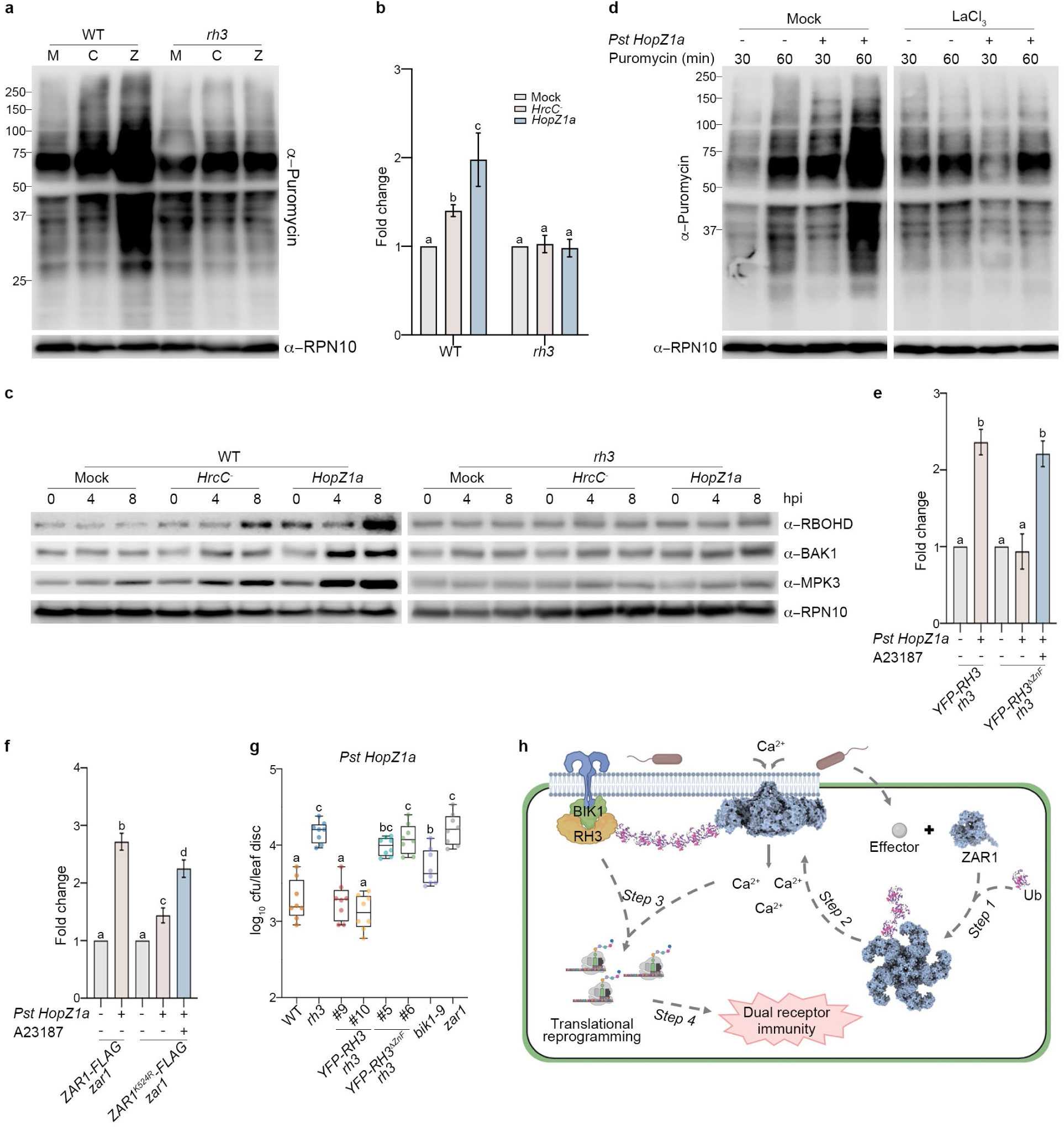
Co-activation of PTI and ETI is underpinned by ubiquitination of ZAR1 and its recognition by RH3 to boost calcium-mediated translational reprogramming. **a,** SUnSET analysis of global translational activity during PTI and ETI. Adult plants were inoculated with H_2_O (M), 5 × 10^6^ CFU/mL *Pst* DC3000 *hrcC⁻* (C), or 5 × 10^6^ CFU/mL *Pst* DC3000 (*HopZ1a*) (Z) for 4 h, followed by infiltration with 50 μM puromycin. Samples were collected 1 h after puromycin treatment. Puromycin incorporation was detected by immunoblotting using an anti-Puromycin antibody, and RPN10 was used as a loading control. **b,** The intensity of puromycin-labeled proteins was quantified, normalized to the mock treatment, and presented as mean ± SD (Tukey’s HSD test following one-way ANOVA; *P* < 0.05, n = 3). **c**, Protein accumulation of PTI components during immune responses. Adult plants were inoculated with the indicated bacterial strain and samples were collected at different time points for western blot analyses. **d**, SUnSET analysis of PTI- and ETI-induced global translational activity in absence or presence of the calcium channel inhibitor LaCl_3_ (0.5 mM). Samples were analyzed as in (**a**). **e**, The calcium ionophore A23187 rescues the translational defect caused by deletion of RH3’s ZnF domain after co-activation of PTI and ETI. Adult plants were inoculated with either H_2_O (Mock) or 5 × 10^6^ CFU/mL *Pst* DC3000 (*HopZ1a*), and with or without 50 μM A23187 for 4 h, followed by infiltration with 50 μM puromycin. Samples were collected 1 h after puromycin treatment. Puromycin incorporation was detected by immunoblotting using an anti-Puromycin antibody, and RPN10 was used as a loading control. The intensity of puromycin-labeled proteins was quantified and normalized to RPN10. The values for *Pst* DC3000 (*HopZ1a*)-treated samples were then normalized to mock-treated samples and presented as mean ± SD (Tukey’s HSD test following one-way ANOVA; *P* < 0.05, n = 3). **f**, The calcium ionophore A23187 partially restores the translational defect caused by mutation of the K524 residue in ZAR1 after co-activation of PTI and ETI. Experiments were performed as in panel **e** above. **g**, Plants were infected with 5 × 10⁵ CFU/mL *Pst* DC3000 (*HopZ1a*) and pathogen growth was evaluated at 3 dpi. **h**, Schematic model of RH3-mediated assembly of a ubiquitinated dual receptor complex. Upon attempted infection, delivery of pathogen effectors induces ZAR1 ubiquitination (step 1), which promotes resistosome assembly (step 2). RH3 specifically binds ubiquitinated ZAR1 and recruits it to cell-surface receptor complexes containing BIK1. Formation of this dual receptor complex potentiates calcium-dependent global translational reprogramming (step 3), thereby establishing dual receptor immunity (step 4).

Calcium plays an important role in eukaryotic protein translation^50^. Given that RH3 recruits the calcium-permeable ZAR1 resistosome to cell-surface receptor complexes, we investigated if calcium is required for the translational boost observed during dual receptor immunity. Indeed, treatment with a calcium channel inhibitor abolished translational upregulation induced by co-activation of PTI and ETI (Fig. 4d). Like *rh3* mutant plants, the *zar1* mutant also exhibited this translational defect, which was rescued by expression of ZAR1-FLAG but not the ZAR1^K524R^ variant (Extended Data Fig. 9a), indicating that ubiquitination of ZAR1 is required for full calcium-dependent translational reprogramming during immunity. Moreover, while *YFP-RH3* expression fully recovered translational activation in the *rh3* mutant, the *YFP-RH3^ΔZnF^*and *YFP-RH3^Q386A^* mutant variants failed to complement this defect (Extended Data Fig. 9b), illustrating that ubiquitin recognition and helicase activities of RH3 are essential for translational reprogramming during co-activation of PTI and ETI. To prove that these collective defects in translational reprogramming are due to loss of calcium influx into the cell, we utilized the calcium ionophore A23187, which elevates intracellular calcium independent of calcium channels. Remarkably, treatment with A23187 restored translational efficiency of the *YFP-RH3^ΔZnF^* and *ZAR1^K524R^-FLAG* mutant variants (Fig. 4, e and f). Together, these data demonstrate that RH3 recruits ubiquitinated ZAR1 to cell-surface receptor complexes to increase the availability of calcium for translational reprogramming.

These findings suggest that RH3 is indispensable for establishing immunity against pathogens that trigger both PTI and ZAR1-dependent ETI. However, loss of *RH3* function had only a moderate effect on immune protein accumulation upon ETI activation alone (Extended Data Fig. 9c), suggesting that PTI activation is required for RH3 to potentiate immune outputs during ETI. To test the biological significance of this, we challenged plants with *Pst* DC3000 strains carrying ZAR1-recognized effectors (*HopZ1a* or *HopO1c*) that trigger co-activation of PTI and ETI, and examined bacterial growth. Despite accumulating elevated levels of ROS upon artificial co-activation of PTI and ETI (Extended Data Fig 9d), the *rh3* mutant exhibited enhanced susceptibility to these avirulent *Pst* DC3000 strains compared to WT and failed to induce a hypersensitive cell death response (HR) (Fig. 4g and Extended Data Fig. 9, e and f). Moreover, while expression of YFP-RH3 fully restored resistance in the *rh3* mutant, the *YFP-RH3^ΔZnF^* mutant failed in this respect. Thus, RH3 and its ability to recognize ZAR1-anchored ubiquitin chains are required for dual receptor immunity. Notably, the *bik1* mutant also displayed enhanced susceptibility compared to WT plants (Fig. 4g), demonstrating further that all components of the RH3-assembled dual receptor complex are required for establishing robust immunity. Furthermore, the *rh3* mutant exhibited impaired induction of translation, defective hypersensitive cell death and compromised immunity against *Pst* DC3000 strains carrying distinct effectors (*i.e.* avrRpt2, HopAR1h, avrB1b) recognized by different plasma membrane-localized NLRs (*i.e.* RPS2, RPS5 and RPM1) (Extended Data Fig. 10), suggesting that RH3 is capable of recruiting different types of dual receptor complexes. Taken together, these findings identify RH3 and associated ubiquitin recognition as a critical new signaling node that integrates both PTI and ETI signals to establish dual receptor immunity.

## Discussion

Cell-surface receptor-mediated PTI and intracellular NLR-mediated ETI form an integrated and mutually reinforcing immune network that enables plants to establish a rapid and robust defense response against diverse pathogens. Yet, it remained unclear how plants integrate signaling by these two types of immune receptors. In this study, we discover that ubiquitination of the NLR receptor, ZAR1, and its recognition by the ZnF-type ubiquitin-binding protein, RH3, act as defining cues for assembly of a dual receptor complex. We propose that RH3-dependent integration of ubiquitinated ZAR1 and cell-surface receptor components into a dual receptor complex enables a cellular influx of calcium that sustains a boost in receptor-induced global translational reprogramming, necessary for establishing immunity (Fig. 4h). Thus, our findings place NLR ubiquitination and ubiquitin recognition at the heart of receptor-mediated immune signaling in plants. The evolutionary conservation of RH3 across plant species (Extended Data Fig. 4a), the rarity of its ZnF domain amongst RNA helicases, its interactions with multiple core PTI and ETI components (Fig. 2), and its requirement for immunity against *Pst* DC3000 strains carrying different effectors recognized by distinct NLRs (Extended Data Fig. 10), suggest that it may facilitate the formation of multiple different dual receptor complexes to establish immunity against adapted pathogens.

The role of ubiquitin in plant dual receptor immunity is reminiscent of innate immunity in humans. Here, activation of the IL-1 receptor (IL-1R) triggers formation of K63- and M1-linked hybrid ubiquitin chains anchored to IRAK1 and TRAF6, two cell-surface receptor complex components. These chains are recognized by the topology-specific ubiquitin readers, TAB2/3 and NEMO, which recruit TAK1 and IKK kinases, respectively, enabling immune signal transduction^51^. This ubiquitin-mediated recruitment of kinase signaling to cell-surface receptor complexes in humans parallels our findings of ubiquitin-mediated recruitment of calcium-permeable NLR resistosome channels to cell-surface receptor complexes in plants. Hence, ubiquitin and the ubiquitin recognition machinery serve as universal spatial organizers of immune signaling in eukaryotic cells.

## Materials and Methods

### Plant materials, growth condition, and plasmid constructions

*Arabidopsis thaliana* and *Nicotiana benthamiana* were germinated under long day condition (16 h light, 8 h dark) at 65% humidity and 22°C with light intensity of 70–100 μmol m^-2^ sec^-1^. Mutant plants used in this study are listed in Extended Data Table 1.

To add epitope tags, the coding sequences of genes were cloned into pENTR/D-TOPO vector or pCR8/GW/TOPO (Invitrogen) and recombined into the destination vector by LR reaction (Invitrogen). Site-directed mutagenesis was performed using the QuickChange Lightning Site-Directed Mutagenesis Kit (Agilent). RH3 and RH3^ΔZnF^ tagged with YFP and RFP were generated by recombining into the pUBQ10:YFP-GW and pUBC-RFP-Dest vector, respectively^52,53^. To generate HA-tagged ZAR1, gene-specific primers with incorporated HA tag at the C-terminus were used to amplify the coding sequence, and recombined into pUBQ10:GW vector^52^. To generate pZAR1:ZAR1-FLAG, the *ZAR1* genomic sequence, including a 1 kb region upstream of the *ZAR1* start codon, was cloned into pEarleyGate 303 vector. The pDEST15 vector was used to generate GST-fusion plasmids. The pMDC7 vector was used to generate the EST:2xFLAG-HopZ1a construct^54^. The primers used in this study are listed in Extended Data Table 2

### Gene expression measurement

Total RNA was extracted as described^55^, cDNA was synthesized by using SuperScript™ II Reverse Transcriptase (Invitrogen) according to the manufacturer’s instructions. qPCR was performed by using PowerUp™ SYBR™ Green Master Mix (Applied Biosystems) on a StepOnePlusTM Real-Time PCR system (Applied Biosystems). Primers used for qPCR are listed in Extended Data Table 2.

For RNA sequencing (RNA-seq) analysis, reads were aligned to the *A. thaliana* TAIR10 genome using Strand NGS. Raw counts were normalized using DESeq with baseline transformation to the median of all samples. Data were then expressed as normalized signal values (i.e., log2(RPKM) where RPKM is read count per kilobase of exon model per million reads) for all statistical tests and plotting. RNA-seq data have been deposited in Array Express at European Molecular Biology Laboratory-European Bioinformatics Institute (EMBL-EBI) under accession codes E-MTAB-15957.

### Protein analysis

Liquid nitrogen frozen plant tissue was ground in protein extraction buffer (50 mM Tris-HCl (pH 7.5), 150 mM NaCl, 5 mM EDTA, 0.1% Triton, 0.2% Nonidet P-40, 50 μg/mL N-p-Tosyl-L-phenylalanine chloromethyl ketone (TPCK), 50 μg/mL Nα-Tosyl-L-lysine chloromethyl ketone hydrochloride (TLCK), 0.6 mM phenylmethylsulfonyl fluoride (PMSF)) unless otherwise stated. Protein extracts were incubated with 1× SDS sample buffer supplemented with 50 mM DTT at 80°C for 10 mins, and separated by SDS-PAGE. All the antibodies used are listed in Extended Data Table 1.

For isolation of plasma membrane fractions, a Minute Plant Plasma Membrane Protein Isolation Kit (Invent Biotechnologies, SM-005-P) was used according to the manufacturer’s instructions. Total protein and plasma membrane proteins were dissolved in 1× SDS sample buffer supplemented with 50 mM DTT at 70°C for 5 mins before separating by SDS-PAGE.

For analysis of ZAR1 ubiquitination level, *N. benthamiana* leaves were infiltrated with *Agrobacterium* strain GV3101 carrying indicated constructs. Leaf tissue was homogenized in extraction buffer (10 mM Tris-Cl (pH 7.5), 150 mM NaCl, 0.5 mM EDTA, 0.5% Nonidet P-40, 10 mM NSC632839, 40 µM MG132, 1 mM DTT, 50 μg/ml TPCK, 50 μg/ml TLCK, 0.6 mM PMSF). Protein extract was incubated with Halo-tag magnetic beads (Promega) immobilized with HALO-Ubiquilin, HALO-NEMO, or HALO-RAD23 to detect total ubiquitination, M1 linkage modified, or K48 linkage modified proteins, respectively. Protein sample was separated by SDS-PAGE, ubiquitination level of ZAR1-HA was detected by immunoblotting with anti-HA. For analysis of ZAR1 ubiquitination levels in Arabidopsis, adult plants expressing *pZAR1:ZAR1-FLAG* or *pZAR1:ZAR1^K524^-FLAG* were used for immunoprecipitation as above, but with anti-FLAG M2 Magnetic Beads (Millipore). Ubiquitinated proteins were detected using anti-Ub (P4D1, Santa Cruz Biotechnology), K48-linked ubiquitin chains were detected using anti-K48 (Cell Signaling), and ZAR1-FLAG was detected using anti-FLAG (Sigma).

For analysis of oligomerization of ZAR1, *N. benthamiana* leaves were infiltrated with *Agrobacterium* strain GV3101 carrying indicated constructs and samples were collected at 36 hours post-infiltration (hpi). Leave tissue was ground in the extraction buffer containing 50 mM Tris-HCl (pH 7.5), 50 mM NaCl, 5 mM MgCl_2_, 10% glycerol, 1 mM DTT, 0.5% Triton, 50 μg/ml TPCK, 50 μg/ml TLCK, 0.6 mM PMSF. Blue native-PAGE was performed as previously described^56^.

For puromycin incorporation assay, adult plants were inoculated with 10 mM MgCl_2_ (mock) or 10 mM MgCl_2_ containing different *Pst* DC3000 strains at OD600 = 0.02. For calcium channel inhibitor studies, 0.5 mM LaCl_3_ was inoculated simultaneously with pathogen infection. For calcium ionophore treatment, 50 µM A23187 (Sigma) was infiltrated into leaves simultaneously with the pathogen. Leaves were then infiltrated with 50 μM of puromycin dihydrochloride (Sigma) at 4 hpi, and samples collected at indicated time points. Protein was extracted as described above and puromycin incorporation was examined by western blotting using anti-puromycin (Merck).

For detecting accumulation of PTI signaling components, adult plant leaves were inoculated with 10 mM MgCl_2_ (mock) or 10 mM MgCl_2_ containing different *Pst* DC3000 strains at OD600 = 0.02. Plant tissue was ground in extraction buffer containing 50 mM Tris–HCl (pH 7.5), 150 mM NaCl, 10% glycerol, 1 mM DTT, 0.5% Nonidet P-40, 1% Phosphatase Inhibitor Cocktail 3, 50 μg/ml TPCK, 50 μg/ml TLCK, 0.6 mM PMSF, 1× SDS sample buffer. Samples were incubated at 70°C for 5 mins before separating by SDS-PAGE.

### Protein-protein interaction assays

For analysis of interaction between ubiquitin-binding proteins and ubiquitin chains, the expression of GST-tagged proteins was induced in *E. coli* BL21 (DE3) cells. Cells were collected and lysed in extraction buffer (1× PBS, 1× Bugbuster (Millipore), 1 mg/ml lysome, 1% Triton, 12.5 U/ml benzonase nuclease, 50 μg/ml TPCK, 50 μg/ml TLCK, 0.6 mM PMSF). GST-tagged protein was purified using Glutathione Sepharose 4B (GE Healthcare) according to the manufacturer’s instructions. Purified protein was then incubated with 400 μl of interaction buffer (1× PBS, 1% Triton, 1 mg/ml BSA, 10 μM NSC632839), containing 0.25 μg of biotin-tagged tetra-K48 (R&D Systems), biotin-tagged tetra-K63 (R&D Systems), or His-tagged tetra-M1 linear (UBPBio) ubiquitin chains, for 1 h in cold room. Beads were washed with interaction buffer, and resuspend in SDS sample buffer containing 50 mM DTT before incubating for 10 mins at 80°C. GST-tagged protein was detected by immunoblotting with anti-GST (Sigma-Aldrich), biotin-tagged tetra-K48 and tetra-K63 chains were detected by using anti-biotin (Cell Signalling Technology), His-tagged tetra-M1 linear chains were detected by using an anti-ubiquitin antibody (anti-ubiquitinylated proteins clone FK2, Merck).

For detecting protein interactions in *N. benthamiana*, leaves were infiltrated with *Agrobacterium* strain GV3101 carrying indicated constructs, samples were collected at 3 days post-infiltration (dpi). Leaf tissue was ground in extraction buffer (10 mM Tris-Cl (pH 7.5), 150 mM NaCl, 0.5 mM EDTA, 0.5 % Nonidet P-40, 50 μg/ml TPCK, 50 μg/ml TLCK, 0.6 mM PMSF). GFP-tagged protein was purified using GFP-Trap agarose (ChromoTek) according to the manufacturer’s instructions. Samples were heated at 80°C for 10 mins in SDS sample buffer supplemented with 50 mM DTT before protein separation by SDS-PAGE.

### Mass spectrometric analysis

For detecting ubiquitin chain-binding proteins, adult plants were treated with 0.5 mM SA or inoculated with *Psm* ES4326 for 24 h. Leaf tissue was ground in extraction buffer (10 mM Tris-Cl (pH 7.5), 150 mM NaCl, 0.5 mM EDTA, 0.5 % Nonidet P-40, 10 μM NSC632839, 40 μM MG132, 50 μg/ml TPCK, 50 μg/ml TLCK, 0.6 mM PMSF). Protein extract was incubated overnight with agarose beads coupled with linear M1 or K48 tetra-ubiquitin chains, or HisPur cobalt resin coupled with His-tagged K63 ubiquitin chains of varying length (Ub_2_-Ub_7_). Resin-bound monoubiquitin was used as negative controls for M1- and K48-linked chains, while empty resin beads were used as a negative control for K63-linked chains. Beads were then washed 3 times with extraction buffer.

For detecting interactors of RH3, leaf tissue from adult plants expressing *pUBQ10:YFP-RH3* was ground in extraction buffer (10 mM Tris-Cl (pH 7.5), 150 mM NaCl, 0.5 mM EDTA, 0.5 % Nonidet P-40, 10 μM NSC632839, 40 μM MG132, 50 μg/ml TPCK, 50 μg/ml TLCK, 0.6 mM PMSF). YFP-RH3 was purified using GFP-Trap according to the manufacturer’s instructions.

For Liquid Chromatography-Mass Spectrometry (LC-MS), samples were digested in a buffer containing 2 M Urea, 50 mM Tris-HCl, 1 mM DTT and 0.1 μg/μl LC-MS grade trypsin. The digestion buffer was incubated with the beads for 30 mins at 37°C, after which it was removed and the digestion was continued overnight at 37°C. Five microliters of the resuspended peptides were analyzed by reversed-phase nano–LC-MS/MS using a nano-Ultimate 3000 LC system and a Lumos Fusion mass spectrometer (Thermo Fisher Scientific). The flow rate was set to 400 nl/min. Peptides were loaded onto a self-packed analytical column (uChrom 1.6, 0.075 mm × 25 cm) and separated using a 67-min gradient buffer A (2% acetonitrile, 0.5% acetic acid) and buffer B (80% acetonitrile, 0.5% acetic acid) as follows: 0 - 16 mins, 2% buffer B; 16 - 56 mins, 3 - 35% buffer B; 56 - 62 mins, 99% buffer B; and 62 - 67 mins, 2% buffer B. Full-scan spectra recording in the Orbitrap was in the range of m/z 350 to m/z 1,400 (resolution: 2.4×10^5^; AGC: 7.5×10^5^ ions). MS2 was performed in the ion trap with an isolation window of 0.7, an AGC of 2×10^4^, an HCD collision energy of 28, rapid scan rate, a scan range of 145 to 1,450 m/z, 50 ms maximum injection time, and an overall cycle time of 1 s. Data were analyzed using FragPipe with the LFQ-MBR preset workflow. The mass spectrometry proteomics data have been deposited to the ProteomeXchange Consortium via the PRIDE partner repository with the dataset identifiers PXD071135, PXD071198, PXD074126, and PXD081809.

### Pathogen growth assays

*P. syringae* strains were grown in LB media supplemented with 10 mM MgCl_2_ and either 100 μg/ml streptomycin (*Psm ES4326*), 50 μg/ml rifampicin (*Pst* DC3000 *hrcC^-^*), or 50 μg/ml rifampicin and 50 μg/ml kanamycin (*Pst* DC3000 *avrRpt2* and *Pst* DC3000 *HopZ1a*). Cultures were grown overnight then centrifuged at 4,000 rpm for 10 mins. Cells were resuspended in 10 mM MgCl_2_ and absorbance was measured at 600 nm before necessary dilutions were made to adjust concentrations to those indicated in figure legends. Plants were infected by pressure infiltration with a syringe through the abaxial leaf surface. For measurement of bacterial growth, a single leaf disc (0.5 cm in diameter) per plant was cut from infected leaves at the stated dpi and ground in 10 mM MgCl_2_. Serial dilutions were plated on LB supplemented with 10 mM MgCl_2_ and appropriate antibiotics, and colonies were counted after 3 days of incubation at 30°C.

### ROS burst assay

Leaf discs from 4-week-old plants expressing *EST:2xFLAG-HopZ1a* were floated on water overnight, then treated with H₂O or 50 µM β-estradiol for 8 h. Elicitation solution (5 μg/mL HRP, 0.2 μM luminol, with or without 1 μM flg22) was added, and luminescence was recorded every 1 min with a 0.3 s signal integration time, using a Tecan Infinite M200 PRO plate reader.

### Confocal microscopy

Leaves of 4-week-old *A. thaliana* expressing *pUBQ10:YFP-RH3* were used for analysis of subcellular localization of RH3. For the bimolecular fluorescence complementation (BiFC) assay, the split-YFP system was used for agroinfiltration of *N. benthamiana* leaves^57^. Samples were collected after 3 days of infiltration. Leaves were stained with 40 µM FM4-64 for 10 mins and imaged using a Leica SP8 confocal microscope.

### Statistical analysis

Statistical analyses for RNA-seq data were performed using Strand NGS. Comparisons between two groups were assessed using an unpaired two-tailed Student’s *t*-test. For comparisons involving three or more groups, one-way analysis of variance (ANOVA) was performed using IBM SPSS Statistics v.27. For pathogen growth box plots, each datapoint corresponds to a biological replicate and plots show median (horizontal line), 25%–75% percentiles (box) and min to max values (whiskers). Letters indicate statistically significant differences between samples (Tukey’s HSD test following one-way ANOVA; *P* < 0.05, n = 8). Graphs were generated with GraphPad.

## Supporting information

Ubiquitin binding proteins in response to Psm

Ubiquitin binding proteins in response to SA treatment

The ZnF-dependent interactome of RH3

Pathogen-responsive RH3 interactors

Immune-responsive diGly sites

Differentially expressed genes during PTI, PTI+ETI, and RH3-regulated genes

## Data availability

All data are available in the main text or the Supplementary Materials. The mass spectrometry proteomics data have been deposited to the ProteomeXchange Consortium via the PRIDE partner repository with the dataset identifiers PXD071135, PXD071198, PXD074126, and PXD081809. RNA-Seq data have been deposited at ArrayExpress under accession number E-MTAB-15957. Materials can be provided by S.H.S. pending scientific review and a completed material transfer agreement. The data that support the findings of this study are available from the corresponding author S.H.S. upon reasonable request.

## Acknowledgments

We thank W. Ma (The Sainsbury Laboratory) for providing the pEarlygate100:3xFLAG-HopZ1a and pUCP20tk:HopZ1a-HA constructs, C. Zipfel (University of Zurich) for providing the *bik1-9* mutant, L. Jiang (The Chinese University of Hong Kong) for providing the *FREE1-RNAi* line, Z. Wu (Southern University of Science and Technology of China) for providing the *rz-1b rz-1c* mutant, and P. Lopez-Calcagno (Newcastle University) for providing the *cp12* mutants. We also thank Dr. S. Haupt and staff of the Plant Growth Facilities (University of Edinburgh). We are grateful to H.-K. Ahn (University of Edinburgh) and K. van Wijk (Cornell University) for valuable discussions.

## Author contributions

Conceptualization: Z.W., B.O.-P, S.H.S.; Investigation: Z.W., T.M., J.S., C.P., A.K., C.S., B.O.-P.; Funding acquisition: S.H.S.; Project administration: H.G., S.H.S.; Writing: Z.W., B.O.-P., S.H.S.

## Supplementary Materials

Extended Data Figs. 1 to 10

Extended Data Tables 1 to 2

References (*1*–*4*)

Supplementary Data 1 to 6

