## Supplementary material for "Ubiquitin recognition integrates plant cell-surface and intracellular receptor immunity": Supplemenary Materials & Figures

**The PDF file includes:**

Extended Data Figs. 1 to 10
Extended Data Tables 1 to 2
References (1–4)

**Other Supplementary Materials for this manuscript include the following:**

Supplementary Data 1 to 6

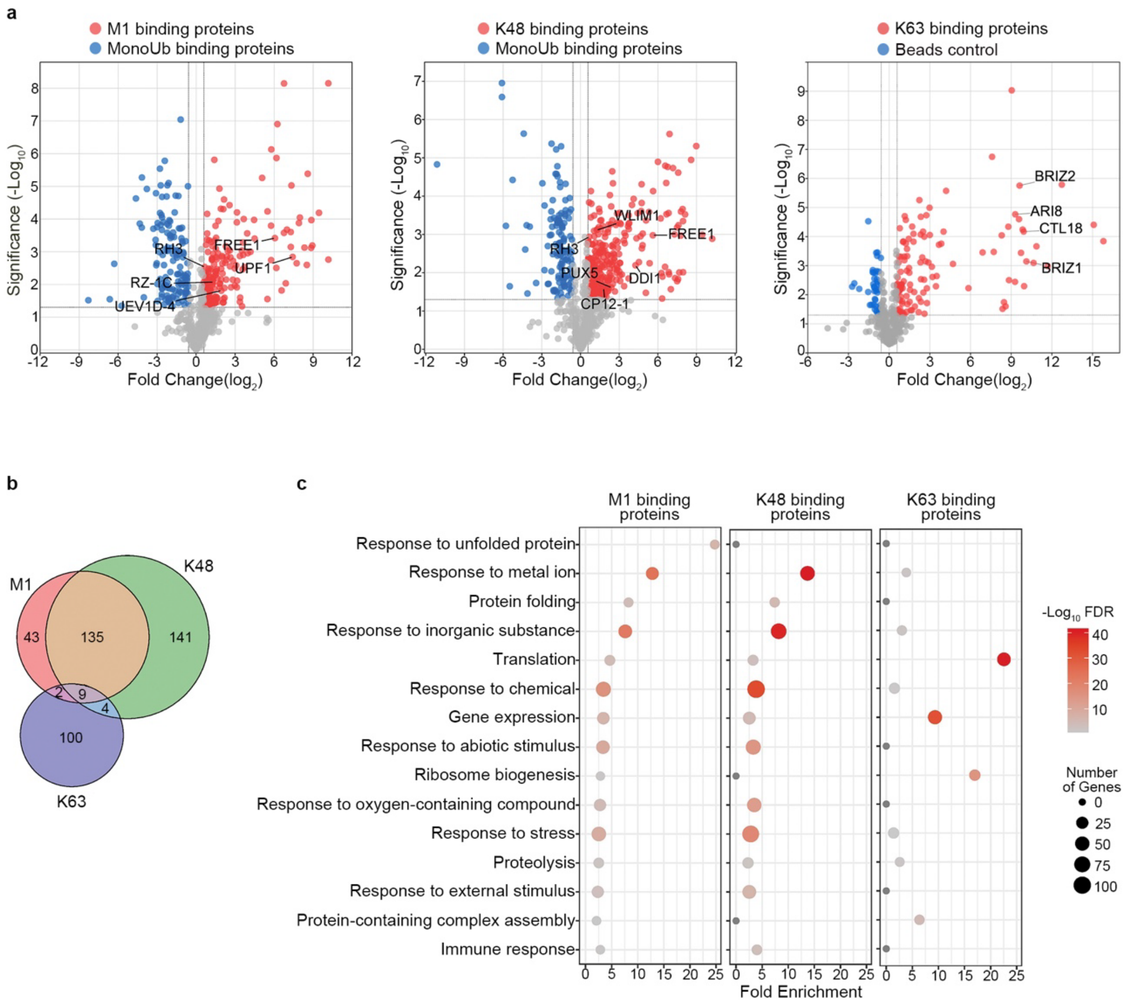


**Extended Data Fig. 1. Identification of SA-responsive M1-, K48- and K63-specific ubiquitin binding proteins. a**, Volcano plots of ubiquitin linkage-specific ubiquitin-binding interactomes. Adult WT plants were treated with 0.5 mM SA for 24 h and binding proteins pulled down with indicated topologies. Monoubiquitin was used as a negative control for M1- and K48-linkage interactomes, and empty beads were used as a negative control for the K63-linkage interactome. Dashed lines indicate thresholds of 0.05 for *P*-values and 1.5 for fold changes relative to the negative control. **b**, Venn diagram showing the overlap between M1-, K48-, and K63-linked ubiquitin chain interactomes. **c**, GO term analysis of linkage-specific interactomes.

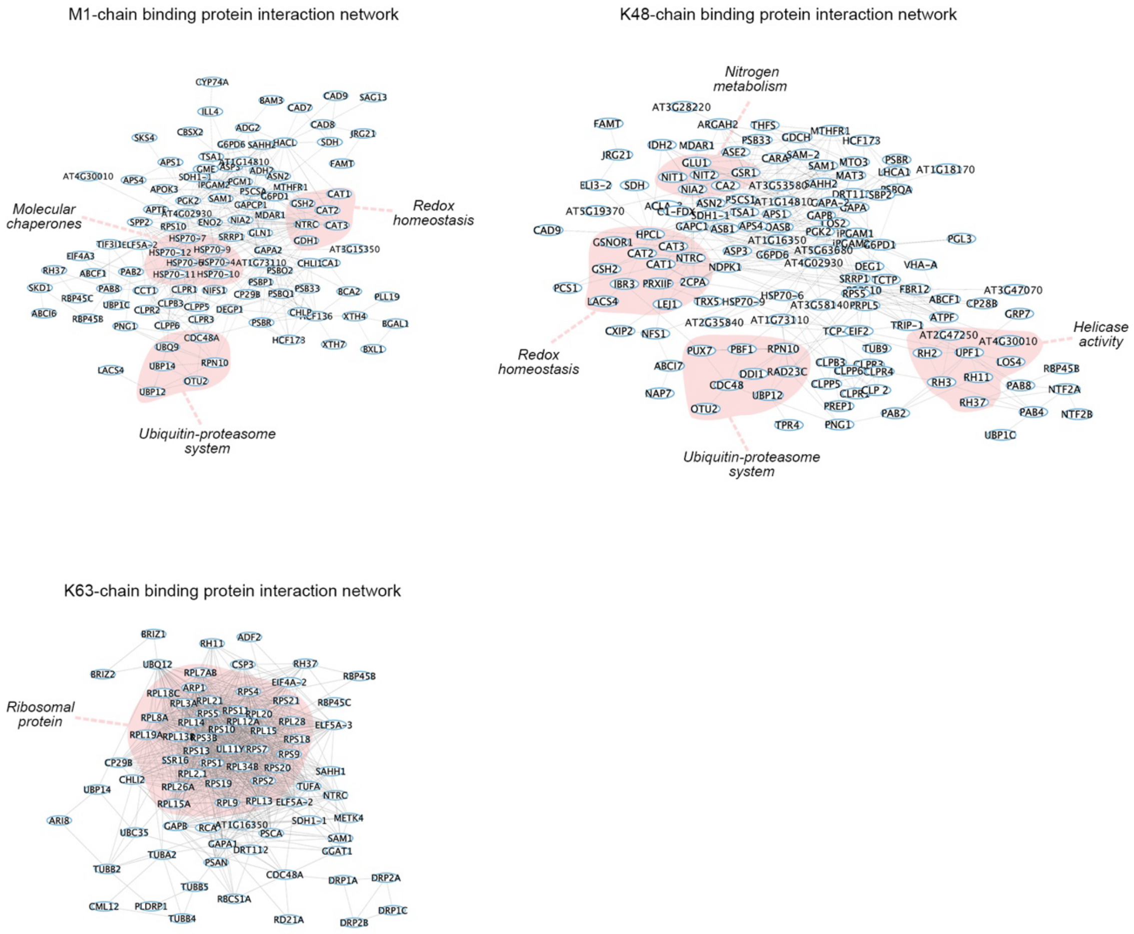


**Extended Data Fig. 2. Protein interaction network of linkage-specific interactomes in response to pathogens.** Ubiquitin topology-specific interactors were analyzed using the STRING database to generate protein-protein interaction networks, which were subsequently visualized and annotated in Cytoscape.

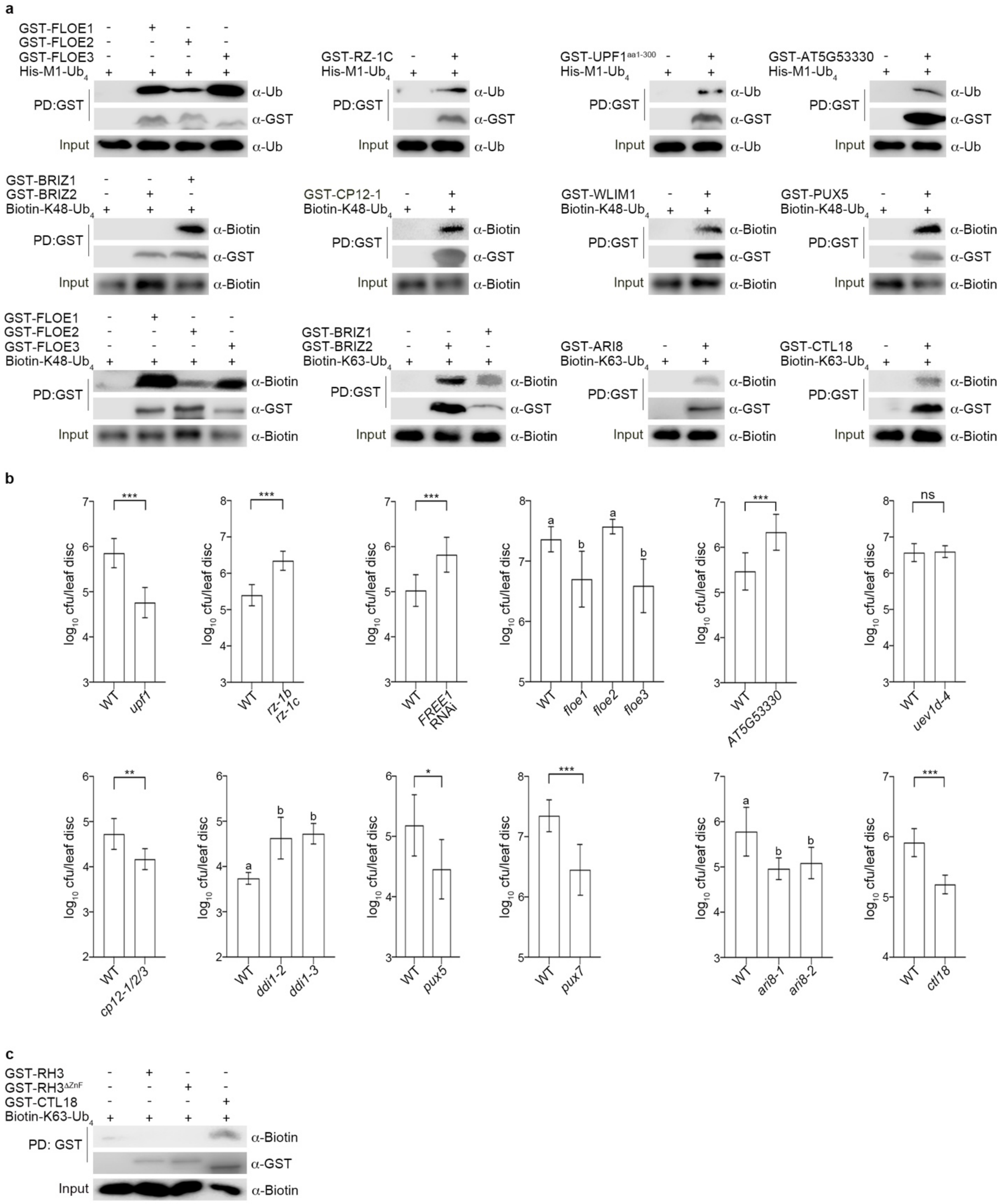


**Extended Data Fig. 3. Ubiquitin-binding proteins are required for immune responses. a**, Ubiquitin-binding proteins (UBPs) physically interact with ubiquitin chains *in vitro*. Recombinant GST-tagged UBPs were purified from *E. coli*, and incubated with M1-, K48-, or K63-linked ubiquitin chains. GST-tagged proteins were detected with anti-GST, His-M1-$Ub_4$ with anti-ubiquitin, and Biotin-K48-$Ub_4$ or Biotin-K63-$Ub_4$ with anti-biotin antibodies. **b**, Loss-of-function mutants of UBPs display compromised disease-responsive phenotypes. Adult plants were inoculated with 5 × $10^5$ CFU/mL *Psm* ES4326, and pathogen growth was evaluated at 3 dpi. Data are presented as mean ± SD (n = 8). Statistical analyses were performed using Student's t tests (**P* < 0.05, ***P* < 0.01, ****P* < 0.001, ns: not significant) or Tukey's HSD post hoc ANOVA tests (different letters indicate significant differences with *P* < 0.05). **c**, RH3 does not interact with K63-linked ubiquitin chains. Experiment was performed as described in **a**. The K63-binding CTL18 protein was used a positive control.

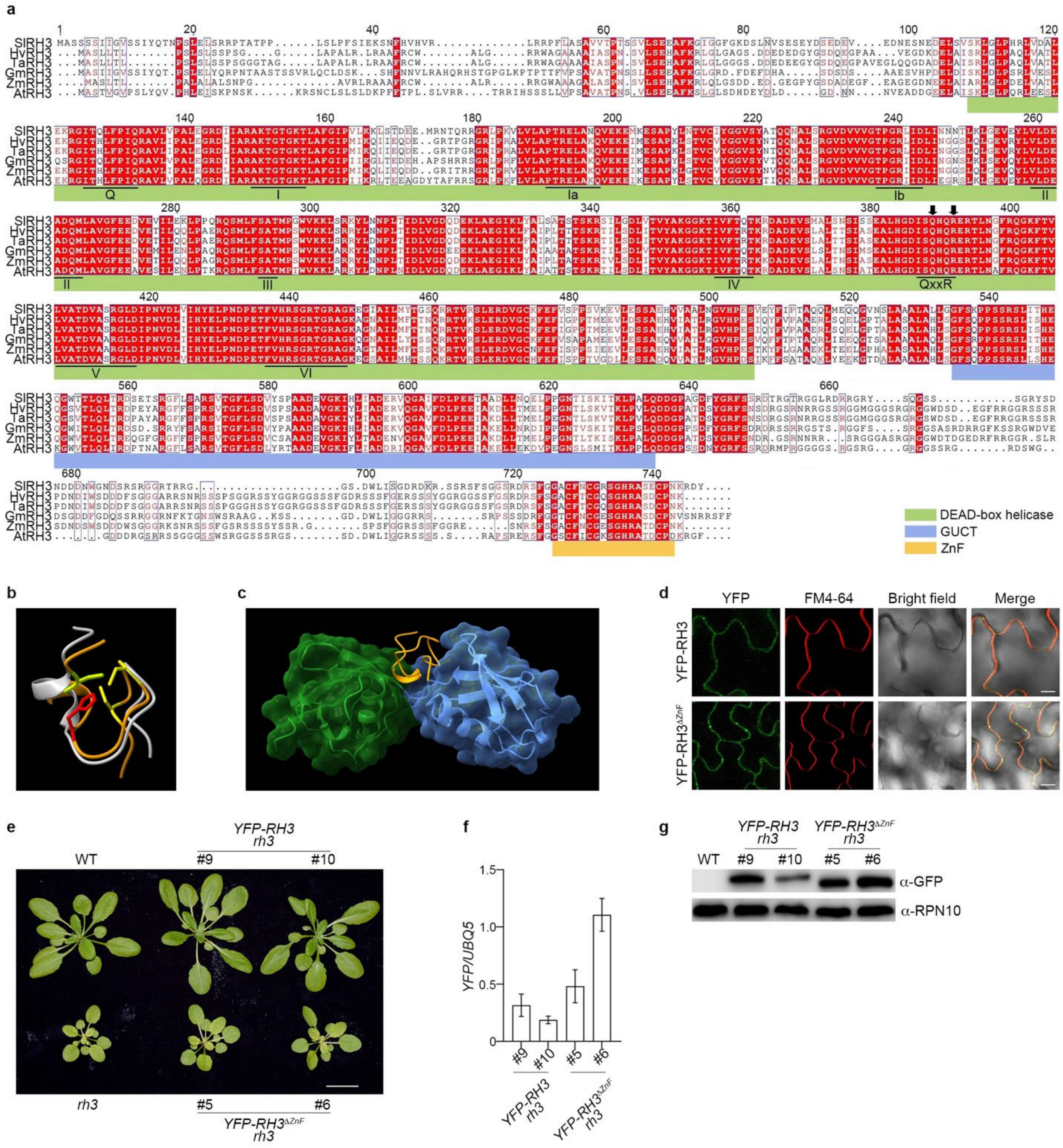


**Extended Data Fig. 4. Characterization of YFP-RH3 transgenic lines. a,** Protein sequence alignment of RH3 orthologs across diverse plant species. Protein sequence alignment of *Solanum lycopersicum* RH3 (XP_004244948), *Hordeum vulgare* RH3 (KAE8793972), *Triticum aestivum* RH3 (XP_044400040), *Glycine max* RH3 (XP_003521635), and *Zea mays* RH3a (PWZ53621). Sequences were aligned using ESPript 3.0. The DEAD-box helicase, GUCT, and ZnF domains are marked with green, blue, and yellow bars, respectively. Black arrows indicate substitution sites within the helicase domain. **b**, The C-terminus of RH3 is predicted to form a CCHC-type ZnF domain. RH3-ZnF (orange ribbon) was modelled onto the its closest predicted structural homologue, the second ZnF domain of the human TUT7 terminal uridylyltransferase (white ribbon). The $Zn^{2+}$-coordinating Cys and His side chains of RH3 are shown in yellow and red, respectively. **c,** AlphaFold Multimer predicts that the RH3-ZnF domain recognizes and bends the interface of M1-linked diubiquitin. **d,** Subcellular localization of RH3 and $RH3^{\Delta ZnF}$ in Arabidopsis. RH3 and $RH3^{\Delta ZnF}$ localize to the plasma membrane. Scale bar = 10 μm. **e**, Developmental phenotypes of adult plants. Scale bar = 2 cm. **f**, Relative mRNA levels of the transgene normalized to *UBQ5* in independent transgenic lines. Data represent mean ± SD (n = 3). **g**, Protein accumulation was detected by immunoblotting with anti-GFP antibody. RPN10 served as a loading control.

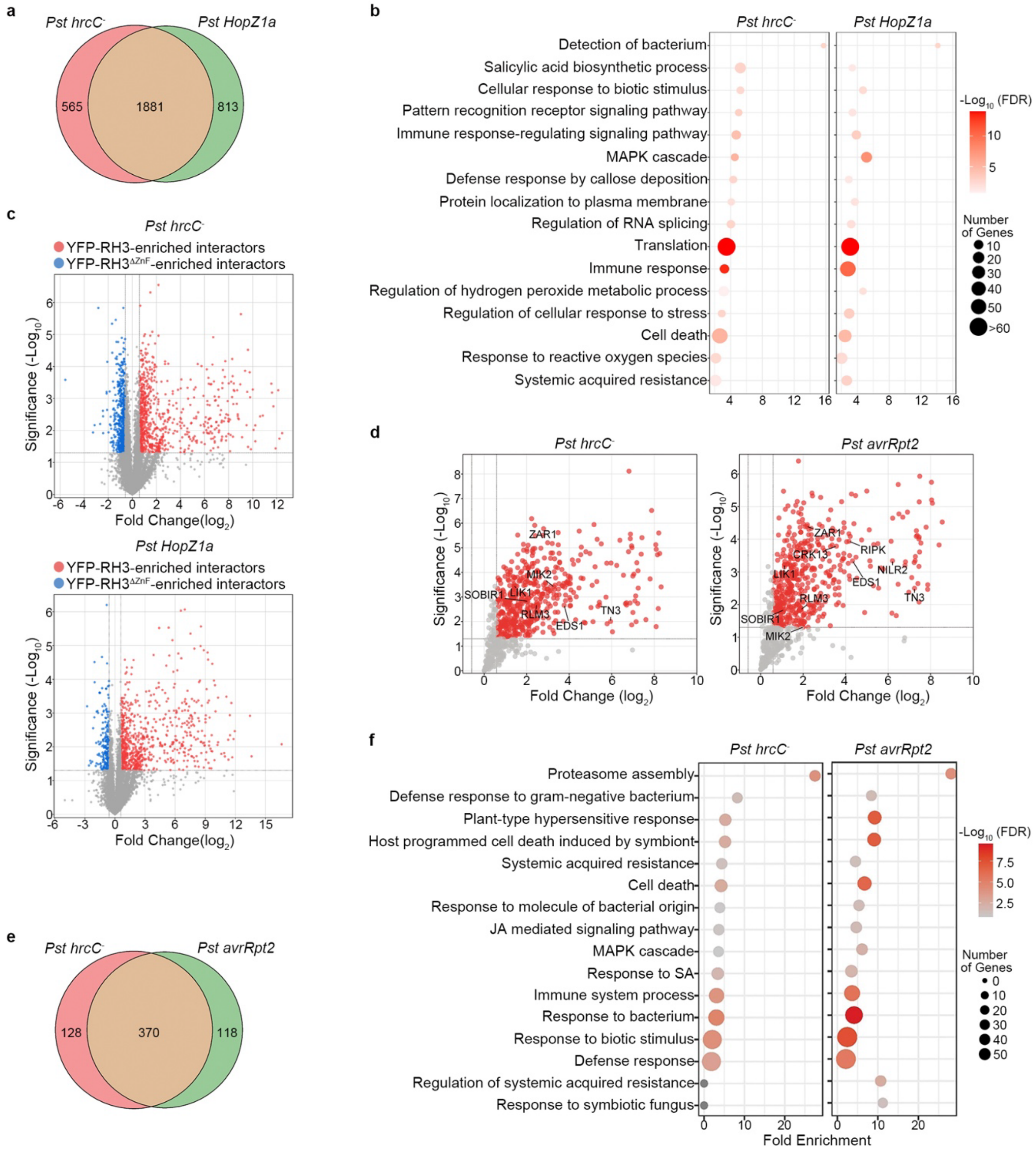


**Extended Data Fig. 5. Identification of RH3 interactors during PTI and PTI + ETI responses.** **a** and **b**, Venn diagram (a) and GO term analysis (b) of RH3 interactors during PTI (*Pst hrcC*$^{-}$) and PTI + ETI (*Pst HopZ1a*). **c**, Deletion of the ZnF domain markedly diminished the RH3 interactome. Upon inoculation with *Pst* DC3000 *hrcC*$^{-}$, 818 and 593 proteins were significantly enriched in the YFP-RH3 and YFP-RH3$^{\Delta ZnF}$ interactomes, respectively. Following inoculation with *Pst* DC3000 (*HopZ1a*), 818 and 227 proteins were significantly enriched in the YFP-RH3 and YFP-RH3$^{\Delta ZnF}$ interactomes, respectively. **d**, Volcano plots of RH3 interactors during PTI (*Pst hrcC*$^{-}$) and PTI + ETI (*Pst avrRpt2*). Plants expressing *YFP-RH3* variants were inoculated with pathogen strains at a concentration of $5 \times 10^6$ CFU/mL and samples were collected at 8 hpi.

Protein complexes were isolated using GFP-Trap magnetic agarose. **e** and **f**, Venn diagram (e) and GO term analysis (f) of RH3 interactors during PTI (*Pst hrcC* $^{-}$) and PTI + ETI (*Pst avrRpt2*).

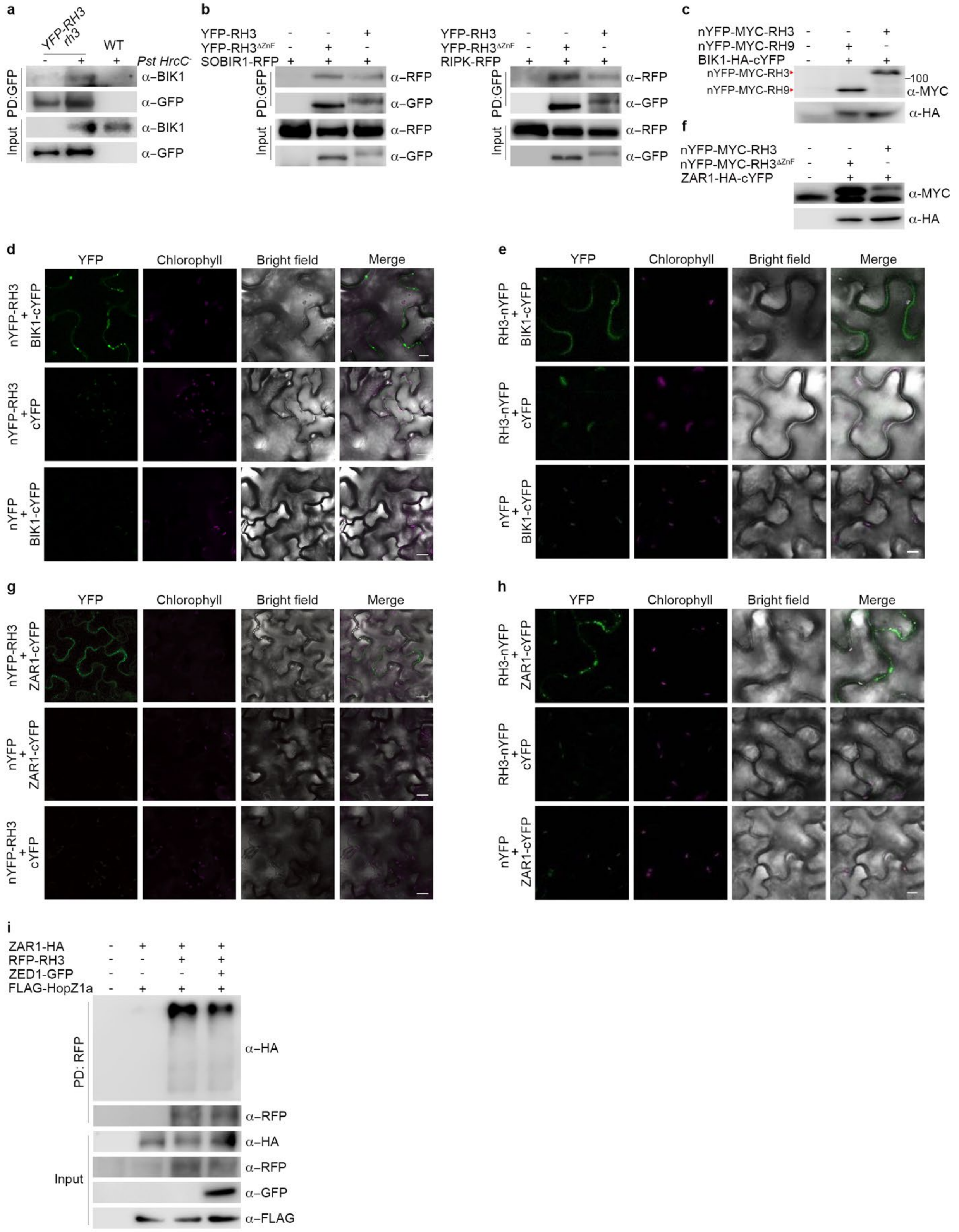
a
YFP-RH3
rh3
WT
Pst HrcC-
PD:GFP
Input
α–BIK1
α–GFP
b
YFP-RH3
YFP-RH3ΔZnF
SOBIR1-RFP
RIPK-RFP
α–RFP
α–GFP
c
nYFP-MYC-RH3
nYFP-MYC-RH9
BIK1-HA-cYFP
100
α–MYC
α–HA
f
nYFP-MYC-RH3ΔZnF
ZAR1-HA-cYFP
d
e
g
h
YFP
Chlorophyll
Bright field
Merge
nYFP-RH3 + BIK1-cYFP
nYFP-RH3 + cYFP
nYFP + BIK1-cYFP
RH3-nYFP + BIK1-cYFP
RH3-nYFP + cYFP
nYFP-RH3 + ZAR1-cYFP
nYFP + ZAR1-cYFP
RH3-nYFP + ZAR1-cYFP
i
ZAR1-HA
RFP-RH3
ZED1-GFP
FLAG-HopZ1a
PD: RFP
Input
α–HA
α–RFP
α–GFP
α–FLAG

**Extended Data Fig. 6. RH3 interacts with both PRR and NLR receptors.** **a**, YFP-RH3 associates with BIK in Arabidopsis. Adult plants were inoculated with $H_2O$ or $5 \times 10^6$ CFU/mL *Pst* DC3000 *hrcC*$^{-}$. YFP-RH3 was isolated using GFP-Trap agarose, followed by immunoblot analysis with anti-BIK1 and anti-GFP antibodies. **b**, RH3 interacts with PTI components. Protein complexes co-expressed in *N. benthamiana* were immunoprecipitated using GFP-Trap magnetic agarose, followed by immunoblotting with indicated antibodies. **c**, Protein expression of the constructs used for BiFC in Fig. 2f was assessed by immunoblot analysis. **d** and **e**, BIK1 interacts with RH3 at the plasma membrane. BiFC assay showing YFP fluorescence occurs at the plasma membrane only when BIK1-cYFP is co-expressed with nYFP-RH3 or RH3-nYFP in *N. benthamiana*. Scale bar = 10 µm. **f**, Protein expression of the constructs used for BiFC in Fig. 3b was assessed by immunoblot analysis. **g and h**, ZAR1 interacts with RH3 at the plasma membrane. BiFC assay showing YFP fluorescence occurs at the plasma membrane only when ZAR1-cYFP and RH3-nYFP are co-expressed in *N. benthamiana*. Scale bar = 10 µm. **i**, The RH3–ZAR1 interaction is not significantly affected by ZED1. Protein complexes co-expressed in *N. benthamiana* were immunoprecipitated using RFP-Trap magnetic agarose, followed by immunoblotting with indicated antibodies.

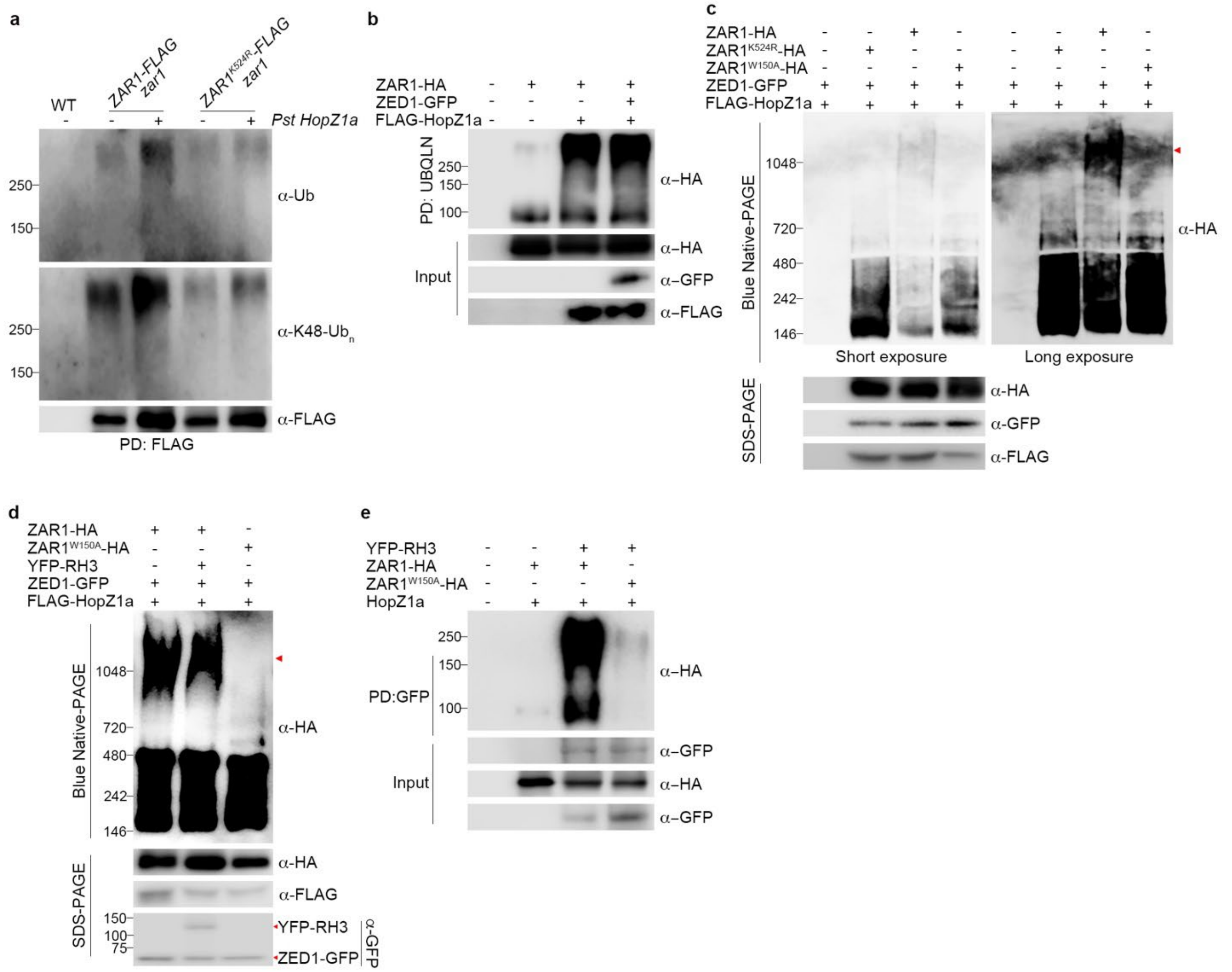


**Extended Data Fig. 7. Ubiquitination promotes ZAR1 resistosome formation independent of ubiquitin recognition by RH3. a**, Bacterial effector HopZ1a induces ZAR1 ubiquitination at K524 in Arabidopsis. ZAR1-FLAG or ZAR1$^{K524R}$-FLAG was immunoprecipitated using anti-FLAG magnetic beads, and ZAR1 ubiquitination was detected by immunoblotting with anti-ubiquitin and anti-K48 linkage-specific antibodies. **b**, ZED1 does not affect ZAR1 ubiquitination. Total ubiquitinated substrates in *N. benthamiana* were pulled down (PD) using recombinant Ubiquilin (UBQLN). ZAR1-HA ubiquitination was detected by immunoblotting with an anti-HA antibody. **c**, BN-PAGE assay shows ubiquitination of K524 promotes HopZ1a-induced oligomerization of ZAR1. Proteins were co-expressed in *N. benthamiana*, and lysate was used for BN-PAGE and SDS-PAGE experiments. Arrowhead indicates the ZAR1 resistosome. **d**, RH3 does not affect HopZ1a-induced ZAR1 oligomerization. Proteins were co-expressed in *N. benthamiana*, and the lysate was used for BN-PAGE and SDS-PAGE analyses. **e**, RH3 pulled down fewer oligomerization-deficient ZAR1$^{W150A}$ mutant compared to wild-type ZAR1 protein in *N. benthamiana*.

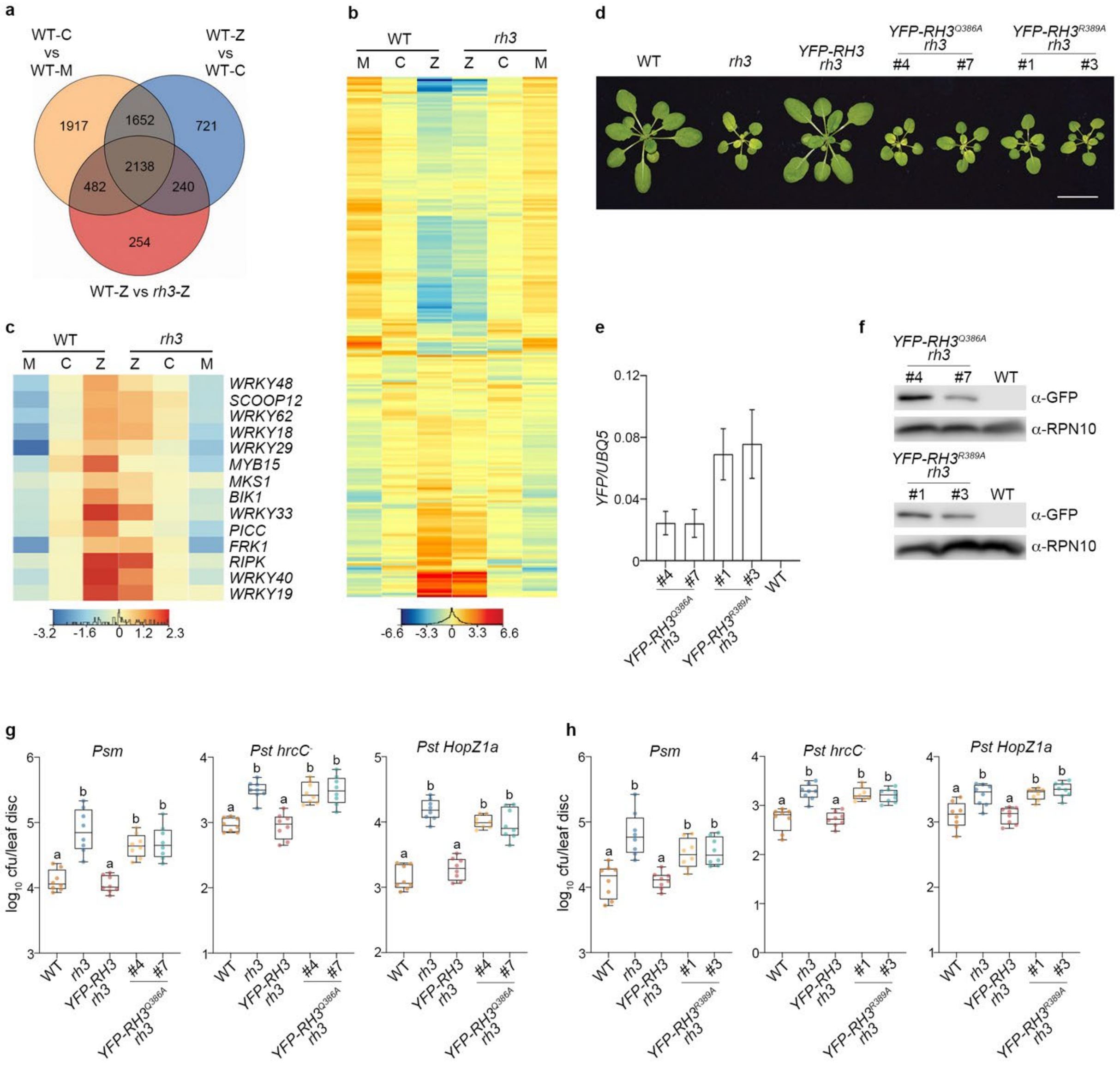


**Extended Data Fig. 8. RH3 has few effects on the transcriptome but its translation-promoting DEAD-box helicase domain is essential for immunity. a-c,** Transcriptional potentiation of PTI during ETI. **a**, Venn diagram showing the overlap between PTI-responsive genes (WT-C vs WT-M), ETI-responsive genes (WT-Z vs WT-C), and RH3-regulated genes (WT-Z vs *rh3*-Z). Genes with a fold change ≥ 1.5 (Benjamini–Hochberg FDR, two-way ANOVA, $P < 0.05$, n = 3) are shown. M: Mock; C: *Pst hrcC*$^{-}$; Z: *Pst avrHopZ1a*. **b**, Heat map showing expression of PTI-responsive genes in WT and the *rh3* mutant. **c**, Expression of subset of immune marker genes analyzed as in (**b**). **d**, Developmental phenotypes of the adult transgenic and mutant plants. Scale bar = 2 cm. **e**, Relative mRNA levels of the transgene normalized to *UBQ5* in independent transgenic lines. Data represent mean ± SD (n = 3). **f**, Protein accumulation detected by immunoblotting with anti-GFP antibody. RPN10 served as a loading control. **g** and **h,** Disease susceptibility assays in YFP-RH3$^{Q386A}$ (d) and YFP-RH3$^{R389A}$ (e) transgenic lines inoculated with indicated bacterial strains at $5 \times 10^5$ CFU/mL and pathogen growth was evaluated at 3 dpi. Letters indicate statistically significant differences between samples (Tukey's HSD test following one-way ANOVA; $P < 0.05$, n = 8).

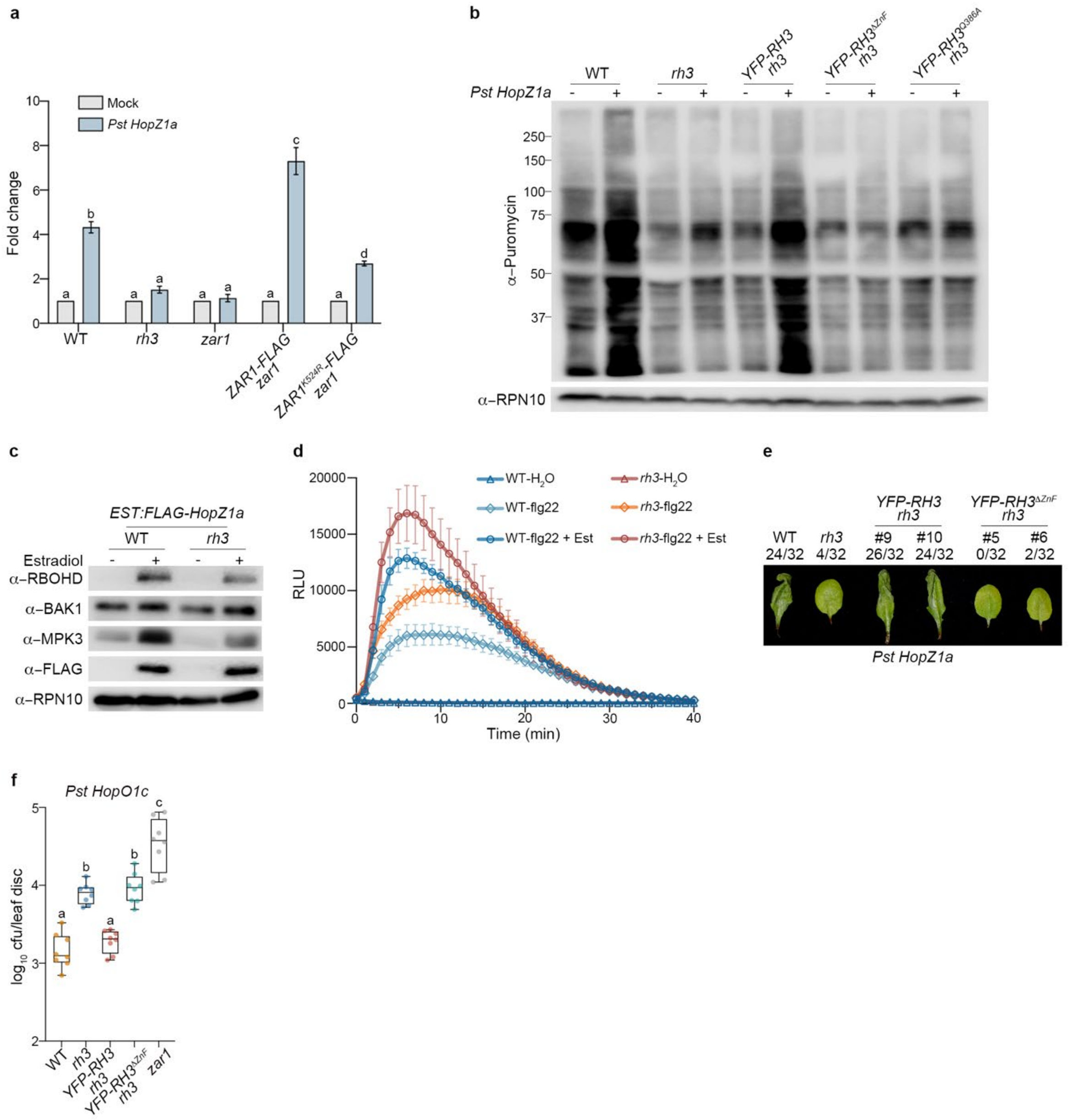


**Extended Data Fig. 9. Co-activation of PTI and ETI is underpinned by ubiquitination of ZAR1 and its recognition by RH3 to boost translational reprogramming. a,** The K524 residue is essential for ZAR1-mediated translational reprogramming during co-activation of PTI and ETI. Adult plants were inoculated with $H_2O$ (Mock) or 5 × $10^6$ CFU/mL *Pst* DC3000 (*HopZ1a*) for 4 h, followed by infiltration with 50 μM puromycin. Samples were collected 1 h after puromycin treatment. Puromycin incorporation was detected by immunoblotting using an anti-Puromycin antibody, and RPN10 was used as a loading control. The intensity of puromycin-labeled proteins was quantified and normalized to RPN10. The values for *Pst* DC3000 (*HopZ1a*)-treated samples were then normalized to mock-treated samples and presented as mean ± SD (Tukey's HSD test following one-way ANOVA; $P < 0.05$, n = 3). **b**, SUnSET analysis of global translational activity in the various *rh3* mutants. Adult plants were inoculated with $H_2O$ or 5 × $10^6$ CFU/mL *Pst* DC3000 (*HopZ1a*) for 4 h, followed by infiltration with 50 μM puromycin. Samples were collected 1 h after puromycin treatment. Puromycin incorporation was detected by immunoblotting using an anti-

Puromycin antibody, and RPN10 was used as a loading control. **c**, Accumulation of PTI components during HopZ1a-induced ETI. Adult plants expressing *Est:FLAG-HopZ1a* were treated with 50 μM estradiol, samples were collected at 8 h post-treatment, and analyzed with antibodies against the indicated proteins. **d**, ROS production during immune responses. Leaves of plants expressing *Est:FLAG-HopZ1a* were treated with $H_2O$, 1 μM flg22, or 1 μM flg22 + 50 μM estradiol, followed by detection of the ROS burst (RLU, relative light units). Data are shown as mean ± SEM (n = 8). **e**, The ZnF domain of RH3 is required for *Pst* DC3000 (*HopZ1a*)-induced HR. Adult plants were inoculated with $H_2O$ (Mock) or $5 \times 10^7$ CFU/mL *Pst* DC3000 (*HopZ1a*). HR phenotypes were scored at 20 h post-infiltration (hpi) by counting the leaves showing cell death relative to the total number of infiltrated leaves. **f**, The ZnF domain of RH3 is essential for establishing resistance against *Pst HopO1c*. Adult plants were inoculated with $5 \times 10^5$ CFU/mL bacteria and pathogen growth was measured 3 days post-infection (dpi). Letters indicate statistically significant differences between samples (Tukey's HSD test following one-way ANOVA; $P < 0.05$, n = 8).

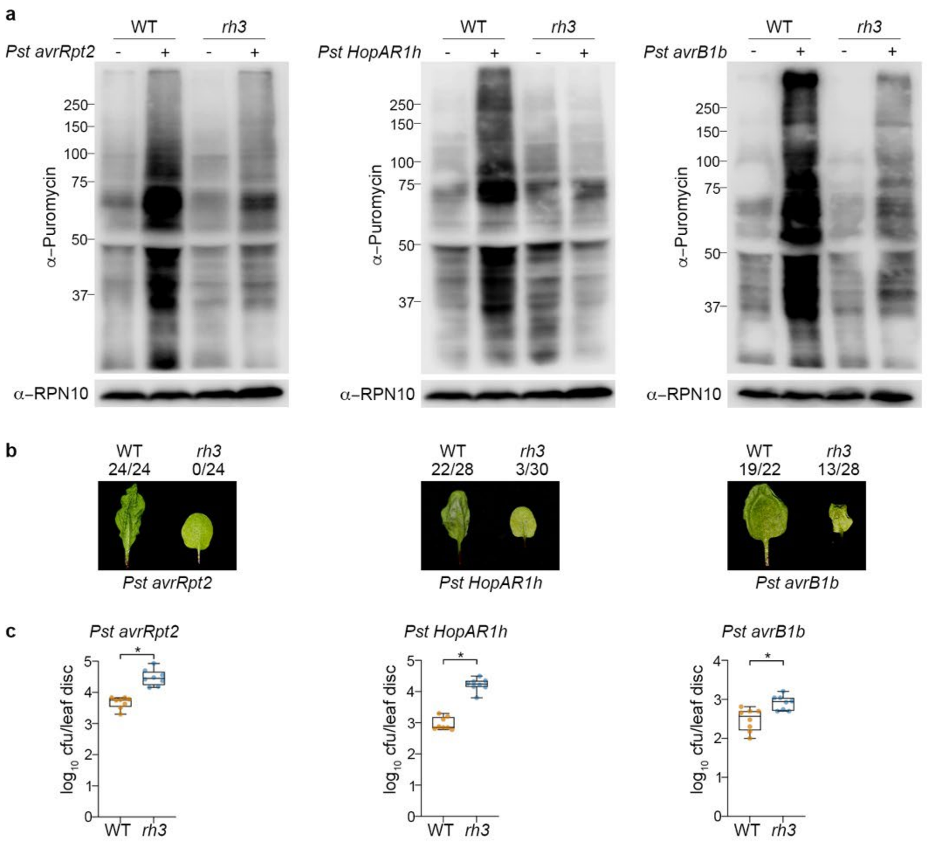


**Extended Data Fig. 10. RH3 is required for translational reprogramming and immunity against distinct NLR receptor-mediated pathways. a**, SUnSET analysis of global translational activity in response to *Pst* DC3000 (*avrRpt2*), *Pst* DC3000 (*HopAR1h*), or *Pst* DC3000 (*avrB1b*). **b**, Adult plants were inoculated with 5 × $10^7$ CFU/mL *Pst* DC3000 (*avrRpt2*), 1.5 × $10^8$ CFU/mL *Pst* DC3000 (*HopAR1h*), 5 × $10^7$ CFU/mL *Pst* DC3000 (*avrB1b*). HR phenotypes were scored at 12 hpi for *Pst* DC3000 (*avrRpt2*) and *Pst* DC3000 (*avrB1b*), and at 24 hpi for *Pst* DC3000 (*HopAR1h*). Data are presented as the number of leaves showing cell death relative to the total infiltrated leaves. **c**, Plants were infected with 5 × $10^5$ CFU/mL *Pst* DC3000 (*avrRpt2*), *Pst* DC3000 (*HopAR1h*), or *Pst* DC3000 (*avrB1b*), and pathogen growth was evaluated at 3 dpi. Letters indicate statistically significant differences between samples (Tukey's HSD test following one-way ANOVA; $P < 0.05$, n = 8).

**Extended Data Table 1. Key resources**

| Reagent type (species) or resource | Designation | Source | Identifier |
|---|---|---|---|
| Antibody | Mouse monoclonal anti-GFP | Roche | Cat# 11814460001 |
| Antibody | Mouse monoclonal anti-Ubiquitin (FK2) | Millipore | Cat# 04–263 |
| Antibody | Mouse monoclonal anti-FLAG M2-Peroxidase (HRP) | Sigma-Aldrich | Cat# A8592 |
| Antibody | Rabbit polyclonal anti-Proteasome 26S S2/PSMD2 | Abcam | Cat# Ab98865 |
| Antibody | Rabbit polyclonal anti-Proteasome 19S S5A/ASF | Abcam | Cat# Ab56851 |
| Antibody | Mouse monoclonal anti-Myc | Sigma-Aldrich | Cat# M5546 |
| Antibody | Mouse monoclonal anti-HA | ThermoFisher | Cat# 26183 |
| Antibody | Mouse monoclonal anti-GST | Sigma-Aldrich | Cat# SAB1305539 |
| Antibody | Anti-Puromycin Antibody, clone 12D10 | Merck | Cat# MABE343 |
| Antibody | Anti-Biotin | Cell Signalling Technology | Cat# 7075 |
| Antibody | Anti-RFP | ChromoTek | Cat# 6G6 |
| Antibody | Anti-RBOHD | Agrisera | Cat# AS15 2962 |
| Antibody | Anti-BAK1 | Agrisera | Cat# AS12 1858 |
| Antibody | Anti-AtMPK3 antibody produced in rabbit | Sigma-Aldrich | Cat# M8318 |
| Antibody | Anti-RH3 | Agrisera | Cat# AS13 2714 |
| Antibody | Anti-PsbO | Agrisera | Cat# AS06 142-33 |
| Antibody | K48-linkage Specific Polyubiquitin | Cell Signaling | Cat# 8081S |
| Antibody | Mouse monoclonal anti-Ubiquitin (P4D1) | Santa Cruz Biotechnology | Cat# sc-8017 |
| Recombinant DNA | pENTR-D-TOPO | Invitrogen | Cat# K240020 |
| Recombinant DNA | pEarleyGate 103 | ABRC | Cat# CD3-685 |
| Recombinant DNA | pEarleyGate 104 | ABRC | Cat# CD3-686 |
| Chemical compound, drug | Recombinant Human Ubiquitin Agarose Protein | R&D Systems | Cat# U-400 |
| Chemical compound, drug | Human Linear Tetra-Ub Non-hydrolyzable Agarose | R&D Systems | Cat# UCN-712 |
| Chemical compound, drug | Human Tetra-Ub Non-hydrolyzable (K48) Agarose | R&D Systems | Cat# UCN-212 |
| Chemical compound, drug | Recombinant Human $His_6$-PolyUb WT Chains (2-7, K63-linked), CF | R&D Systems | Cat# UCH-330 |
| Chemical compound, drug | Recombinant Human Tetra-Ub WT Chains (K48-linked) Biotin | R&D Systems | Cat# UCB-210 |

| | | | |
|---|---|---|---|
| Chemical compound, drug | Recombinant Human Tetra-Ub WT Chains (K63-linked) Biotin | R&D Systems | Cat# UCB-310 |
| Chemical compound, drug | 6xHis-linear $Ub_4$ | UBPBio | Cat# D4210 |
| Chemical compound, drug | MG132 | Cayman Chemical | Cat# 10012628 |
| Chemical compound, drug | Phosphatase Inhibitor Cocktail 3 | Sigma-Aldrich | Cat# P0044 |
| Chemical compound, drug | NSC632839 | Abcam | Cat# ab144599 |
| Chemical compound, drug | Puromycin dihydrochloride | Sigma-Aldrich | Cat# P8833 |
| Chemical compound, drug | GFP-Trap® Agarose | ChromoTek | Cat# gta |
| Chemical compound, drug | Magne® HaloTag® Beads | Promega | Cat# G7282 |
| Chemical compound, drug | Calcium Ionophore A23187 | Sigma-Aldrich | Cat# C7522 |
| Chemical compound, drug | β-Estradiol | Sigma-Aldrich | Cat# E8875 |
| Chemical compound, drug | Horseradish peroxidase | Sigma-Aldrich | Cat# P6782 |
| Chemical compound, drug | Luminol | Sigma-Aldrich | Cat# A8511 |
| *Arabidopsis thaliana* | *upf1* | | SALK_112922 |
| *Arabidopsis thaliana* | *rz-1b rz-1c* | Ref. [1] | SALK_001328 GABI_030b12 |
| *Arabidopsis thaliana* | *rh3-4* | ABRC/NASC | SALK_005920 |
| *Arabidopsis thaliana* | *DEX::RNAi-FREE1* | Ref. [2] | |
| *Arabidopsis thaliana* | *floe1* | ABRC/NASC | SALK_048257 |
| *Arabidopsis thaliana* | *floe2* | ABRC/NASC | SALK_145362 |
| *Arabidopsis thaliana* | *floe3* | ABRC/NASC | SAIL_830_D05 |
| *Arabidopsis thaliana* | *at5g53330* | ABRC/NASC | SALK_067730 |
| *Arabidopsis thaliana* | *uev1d-4* | ABRC/NASC | SALK_064912 |
| *Arabidopsis thaliana* | *cpl12-1 cpl12-2 cpl12-3* | Ref. [3] | |
| *Arabidopsis thaliana* | *ddi1-2* | ABRC/NASC | SAIL_1239_C12 |
| *Arabidopsis thaliana* | *ddi1-3* | ABRC/NASC | SALK_045097 |
| *Arabidopsis thaliana* | *pux5* | ABRC/NASC | SALK_116699 |
| *Arabidopsis thaliana* | *pux7* | ABRC/NASC | WiscDsLox481-484J16 |
| *Arabidopsis thaliana* | *ari8-1* | ABRC/NASC | SALK_058846 |
| *Arabidopsis thaliana* | *ari8-2* | ABRC/NASC | SALK_082904 |
| *Arabidopsis thaliana* | *ctl18* | ABRC/NASC | SAIL_218_F09 |
| *Arabidopsis thaliana* | *zar1-1* | ABRC/NASC | SALK_013297 |
| *Arabidopsis thaliana* | *bik1-9* | Ref. [4] | |

**Extended Data Table 2. List of primers used in this study**

| Name | Purpose | Sequences (5'-3') |
|---|---|---|
| ZAR1-HA | Cloning | F: ATGGTGGACGCTGTTGTAACAG<br>R: TCAagcgtaatctggaacatcgtatgggtaGGTTCTGTGCAATGGTGTTTTC |
| ZAR1$^{K524R}$ | mutagenesis | F: GTTAACTTTGATCTGCCTCTCATCAAAGTTTCCACTAATCCCA<br>R: TGGGATTAGTGGAAACTTTGATGAGAGGCAGATCAAAGTTAAC |
| ZAR1$^{W150A}$ | mutagenesis | F: TCATACACCGGAGAGCTCGCCCGATCAGTTCCATTATC<br>R: GATAATGGAACTGATCGGGCGAGCTCTCCGGTGTATGA |
| RH3 | Cloning | F1: caccATGGCGTCGACGGTAGGAGT<br>R1: CTAAAATCCTCTCTTATCAG |
| RH3$^{\Delta ZnF}$ | Cloning | F: caccATGGCGTCGACGGTAGGAGT<br>R: CTACCAATCATCAGAACTTCCTCT |
| RH3$^{Q386A}$ | mutagenesis | F: GTTCTCTCTCTTTGATGCGCAGATATATCTCCATGAAGTGCTTCG<br>R: CGAAGCACTTCATGGAGATATATCTGCGCATCAAAGAGAGAGAAC |
| RH3$^{R389A}$ | mutagenesis | F: GAAAGCATTGAGTGTTCTCTCTGCTTGATGCTGAGATATATCTCCATG<br>R: CATGGAGATATATCTCAGCATCAAGCAGAGAGAACACTCAATGCTTTC |
| RH3$^{T407A}$ | mutagenesis | F: ACGAGATGCAACATCAGCGGCAACTAATACGGTGA<br>R: TCACCGTATTAGTTGCCGCTGATGTTGCATCTCGT |
| RH3$^{D415A}$ | mutagenesis | F: ATCTACGTTCGGGATGGCAAGTCCACGAGATGC<br>R: GCATCTCGTGGACTTGCCATCCCGAACGTAGAT |
| UPF1$^{aa1\text{-}300}$ | Cloning | F: caccATGGATTCTCAACAGAGCGAT<br>R: tcaTTCAAGGTCTTCAAGAGTAG |
| RZ-1C | Cloning | F: ATGGCTGCAAAAGAAGGTAG<br>R: TTAATAACGGTCAAAAGTGGAC |
| FLOE1 | Cloning | F: caccATGGCGTCTGGATCTTCGGGT<br>R: TCACCATCCTCTGGGAGGTCCT |
| FLOE2 | Cloning | F: caccATGCAATCTTTCGATCTAATA<br>R: CTAACGACCACCAAACCAACCT |
| FLOE3 | Cloning | F: caccATGAATACTTGTCAGTTTATG<br>R: CTACCGACCACCAAAAAAACCT |
| AT5G53330 | Cloning | F: caccATGGATTACGATTACAGAAAC |

| | | |
|---|---|---|
| | | R: TTAGGAAGAACCATGGAGCAA |
| BRIZ1 | Cloning | F: caccATGTTCATCCTCAGAGTTCA<br>R: TTACTTTCTCCGGTTTGACT |
| BRIZ2 | Cloning | F: caccATGAACTCGGCAAGCGTTTC<br>R: CTAACCCTTTCTTCTGTTACT |
| CP12-1 | Cloning | F: ATGACAACCATAGCTGCAGCTG<br>R: TTAATTATCATAAGTACGAC |
| WLIM1 | Cloning | F: ATGGCGTTCGCAGGAACAAC<br>R: TTAAGCAGCGACGACTTTGTC |
| PUX5 | Cloning | F: ATGGCGACGGAGACGAACGAGA<br>R: CTAGAATTTCTGGATGACGAC |
| ARI8 | Cloning | F: caccATGGAAGCTGATGACGATTTC<br>R: TCACCGGCCATGTTCACACA |
| CTL18 | Cloning | F: caccATGCCCATGGAGAACGACAA<br>R: TCAGCTTTGTCCAGAGGTCGA |
| YFP/GFP | qPCR | F: AAGCTGACCCTGAAGTTCATCTGC<br>R: CTTGTAGTTGCCGTCGTCCTTGAA |
| UBQ5 | qPCR | F: CCAAGCCGAAGAAGATCAAG<br>R: ACTCCTTCCTCAAACGCTGA |

**Supplementary Data 1. (separate file)**

Ubiquitin binding proteins in response to *Psm*

**Supplementary Data 2. (separate file)**

Ubiquitin binding proteins in response to SA treatment

**Supplementary Data 3. (separate file)**

The ZnF-dependent interactome of RH3

**Supplementary Data 4. (separate file)**

Pathogen-responsive RH3 interactors

**Supplementary Data 5. (separate file)**

Immune-responsive diGly sites

**Supplementary Data 6. (separate file)**

Differentially expressed genes during PTI, ETI, and RH3-regulated genes
